# Early life adversity increases striatal dopamine D1 receptor density and promotes social alcohol drinking in mice, especially males

**DOI:** 10.1101/2025.11.10.687736

**Authors:** Lucy G. Anderson, Anna Tischer, Roland Bock, Michael Michaelides, Veronica A. Alvarez

## Abstract

The brain’s reward circuitry remains sensitive to experience throughout early life and into adulthood, allowing individuals to adapt to their unique environments. Adverse experiences early in life can increase vulnerability to substance use disorders, likely through alterations to this circuitry. Yet, the precise neurobiological mechanisms by which early life adversity acts are incompletely characterized. In this study, we used a limited bedding and nesting (LBN) paradigm as a translationally relevant model of early life adversity in genetically-identical C57BL/6J mice. After LBN-rearing, we assess the lasting behavioral and neurobiological impacts of this experience in adulthood. In robust sample sizes, our results validated previous findings of increased risk avoidance, enhanced acute response to alcohol, and greater voluntary alcohol drinking in socially-housed LBN-reared mice, especially males. Further, using autoradiography, we found LBN-reared mice had increased striatal D1-like receptor binding, skewing D1-to D2-like receptor balance relative to cross-fostered controls. However, after voluntary alcohol drinking, we found a strong downregulation in D1-like, and some D2-like, receptor binding, negating pre-existing differences in striatal dopamine receptor binding. We posit that via both transcriptional and post-transcriptional mechanisms, LBN-rearing upregulates striatal D1-receptor density and alters risk avoidance and acute alcohol stimulation to promote alcohol drinking among adversity-exposed mice. Together, these findings reveal specific neurobiological mechanisms that promote alcohol consumption following early life adversity and suggest complex interactions between early life adversity, sex-related factors, and dopamine receptor regulation in contributing to alcohol use disorder (AUD) vulnerability.

## INTRODUCTION

Neurons within the brain’s reward circuitry—dopaminergic projections from the ventral tegmental area (VTA) to the nucleus accumbens (NAc) and dorsal striatum—are known to undergo protracted development through adolescence [1–4]. As a result, experiences early in life can change how these neural circuits develop and respond to reward in adulthood [5–11]. This property is key to an individual’s ability to adapt to their environment. Yet, in the case of early life adversity (ELA), such as physical and emotional trauma, neglect, and resource scarcity, experience-induced changes can have maladaptive repercussions in adulthood. Affecting an estimated ∼60% of adults in the United States [12], ELA is known to robustly and dose-dependently confer vulnerability to both mood and substance use disorders (SUDs), including alcohol use disorder (AUD) [13–21]. However, the precise neurobiological mechanisms by which ELA alters reward circuitry to increase AUD vulnerability remain unknown. Furthermore, it is unclear how other factors—including the dimensionality of the adversity (i.e., threat vs deprivation) [8,22,23], sex-related factors [24–28], experiences later in life/cumulative effect [29–32], and the development of comorbid mood disorders [33–37]—may mediate the effects of ELA.

Inbred rodent strains can aid in the isolation of the epigenetic and post-translational mechanisms linking adversity and AUD. In agreement with clinical findings, previous studies have reported altered reward learning and motivation in animal models of ELA [38–41], including increased AUD-like behavior [42–44]. ELA has also been found to alter expression of dopamine D1-[45] and D2-receptors [46] in the NAc, in addition to inducing broader transcriptomic changes across the NAc, VTA, and prefrontal cortex [47]. Despite its established role in flexible decision-making [48,49], the transcriptional effects of ELA on the dorsomedial striatum (DMS) have yet to be investigated. Whether or how transcriptional findings differ from post-transcriptional receptor binding is also unknown.

Striatal dopamine D1-receptor and D2-receptor signaling is known to play a key role in promoting AUD-like behaviors. In line with reports linking increased D1-receptor activation to greater alcohol seeking and consumption in rodents [50–54], we have previously shown that striatal D1-receptor activation is required for alcohol-induced stimulation [55]. In turn, high acute stimulation and low sedation in response to alcohol is robustly linked to clinical AUD vulnerability [56–63]. Imbalance of dopamine D1-to D2-receptors, and broader imbalance between direct (D1-expressing) and indirect (D2-expressing) medium spiny neuron projection pathways within the basal ganglia, has also been linked to SUD vulnerability. We have found that mice with altered D1-to D2-receptor balance show increased baseline risk avoidance and alcohol relief, as well as more punishment-insensitive alcohol drinking [55,64]. This result is consistent with clinical findings of low D2-receptor availability in individuals with AUD [65–70].

Our study sought to understand the impact of ELA on reward-related behavioral and mechanistic adaptations. To induce adversity, we used the limited bedding and nesting (LBN) paradigm, which induces more unpredictable and fragmented maternal care [71–76]. Validating previous findings, we found that LBN-reared mice had greater risk avoidance, enhanced acute response to alcohol, and greater voluntary alcohol drinking in a social setting relative to cross – fostered controls. Differences in drinking were most pronounced among males. Our novel study of striatal D1– and D2-like receptor binding and expression revealed adversity-induced increases to striatal D1-like receptor binding, especially in the NAc. After voluntary alcohol drinking, however, we found robust downregulation of D1-like receptor expression and binding, in both control and LBN-reared mice, which negated pre-existing differences in striatal dopamine receptor binding. Together, our results point towards mediating influences of sex-related factors, risk avoidance, and alcohol on the effects of early life adversity on striatal dopamine receptor density and alcohol drinking behavior.

## MATERIALS AND METHODS

### Animals

Timed-pregnant dams (gestational day 17, n = 33; C57BL6/J background, JAX: 000664) were purchased and rehoused into standard housing upon arrival. A total of 187 pups (Control: n = 42 F, 51 M; LBN: n = 33 F, 61 M; C57BL6/J background, JAX: 000664) were used across all experimentation. Of those, 175 pups (Control: n = 41 F, 49 M, mortality: 3.23%; LBN: 31 F, 54 M, mortality: 9.57%) survived until post-natal day (PND) 60. Cohorts one (n = 18 F, 22 M), two (n = 13 F, 17 M), and three (n = 16 F, 18 M) completed light dark box, open field, repeated elevated zero maze, ethanol-induced locomotion, and social operant alcohol drinking tests prior to sacrifice. Cohorts four (n = 11 F, 14 M) and five (n = 14 F, 32 M) completed light dark box, elevated zero maze, and social operant water drinking tests prior to sacrifice. Pups were aged PND 55-150 at the time of behavioral testing. For all materials and methods, refer to supplementary information for more detail.

### Limited Bedding and Nesting Paradigm

On PND 3, mice were cross-fostered and randomly assigned to LBN or control housing conditions, with equivalent litter sizes (∼6-7 mice) and 1:1 sex ratios. Cages remained undisturbed until PND 11, after which mice returned to standard housing conditions. The LBN conditions used in this study were consistent with that described previously [71,75]. More in supplement.

### Behavior

Paradigms for the light-dark box (LDB), open field (OF), elevated zero maze (EZM) and repeated EZM, and ethanol-induced locomotion tasks used in this study were consistent with previously published lab protocol [64]. More in supplement.

### Social Operant Drinking

The IntelliCage Testing Systems (TSE Systems GmbH, St Louis, MO, USA) were used for automated tracking of social, operant drinking behaviors in a homecage environment. The IntelliCage apparatus consisted of a large cage (Tecniplast 2000, West Chester, PA, USA) equipped with 4 operant corner chambers, each containing two sipper bottles. Up to 16 same-sex mice were housed together per cage, each uniquely identified with a subcutaneously implanted radiofrequency identification (RFID) transponder (1.25 x 7 mm; TSE).

Mice individually entered operant corners via a small tube equipped with a RFID antenna and temperature sensor, both of which required a positive signal for sipper bottle access. Visit, nose poke, and lick data was recorded for each mouse. Animal weight and wellness was checked during weekly cage changes. More in supplement.

### Quantitative Polymerase Chain Reaction

Expression of *Drd1* and *Drd2* mRNA was measured relative to region-specific group averages among alcohol-naïve control mice via the ΔΔCt method using *Actb* as the internal control gene. Methods were consistent with previously published lab protocol [64]. More in supplement.

### Autoradiography

Percent specific binding of [^3^H]raclopride (4 nM, 81.8 Ci/mmol, Revvity, Waltham, MA, USA) and [^3^H]SCH-23390 (2.5 nM, 83.9 Ci/mmol, Revvity) was assessed in thaw-mounted sections and manually analyzed across striatal subregions relative to region-specific group averages among alcohol-naïve, control mice. Methods were consistent with previously published lab protocol [64]. More in supplement.

### Fast-Scan Cyclic Voltammetry

Fast-scan cyclic voltammetry (FSCV) recordings were conducted in the DMS consistent with previously published lab protocol [55,64]. Transients were compared before and after bath application of 1 μM DHßE. More in supplement.

### Statistical Analysis

Data were analyzed and graphed in R using publicly available packages from the tidyverse. Data with repeated measures (e.g., body weight, drinking, locomotion) were analyzed via mixed methods ANOVA (mixed ANOVA) employing a Greenhouse-Geisser sphericity correction. Subsequent pairwise comparisons were made using multiple t-tests with Bonferroni adjustment. Data without repeated measures (e.g., risk avoidance, relative mRNA expression, binding) were analyzed via unpaired t-test or independent measures ANOVA when multiple between-subject factors were assessed. Post-hoc analysis used Tukey’s multiple comparisons test. Correlations between continuous datasets were analyzed via linear regression. All code is available upon request.

## RESULTS

### Body weight is lower in LBN-exposed mice throughout lifespan, especially in males

Following completion of the LBN paradigm (PND 11), male and female LBN-exposed mice weighed less than their control counterparts. This pattern persisted throughout the experiment (Figure 1B), with the exception of female mice on PND 60.

**Figure 1.**
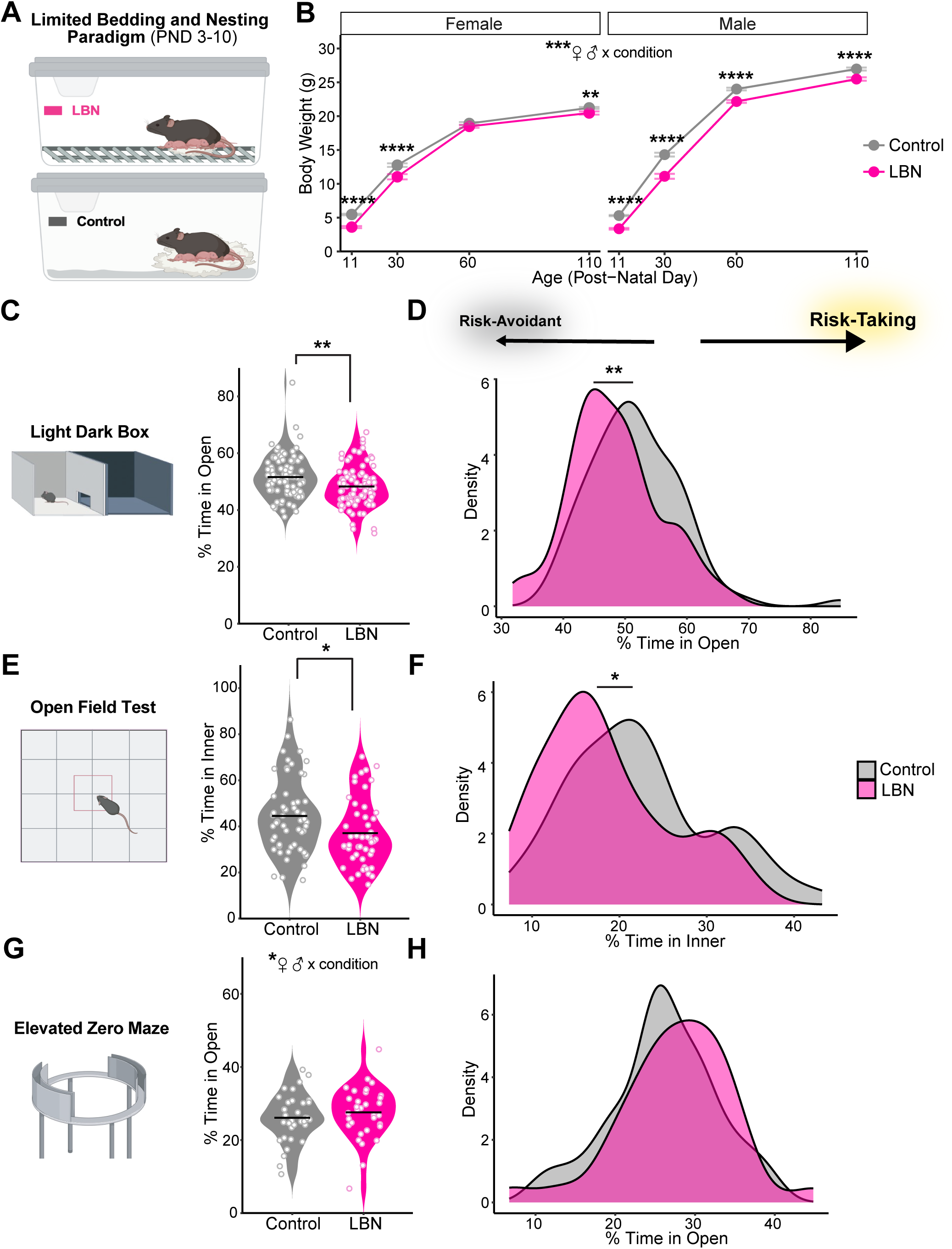
Limited bedding and nesting (LBN) paradigm lowers body weight and increases risk avoidance behavior in C57BL/6J mice. (A) Schematic of LBN (top) and control (bottom) rearing conditions. (B) Body weight (g) of female (left) and male (right) mice reared under LBN (15 litters; n = 40 F, 61 M) and control (16 litters; n = 47 F, 58 M) control conditions (Mixed ANOVA, condition: F(1,164) = 101.68, p = 6.59e-19; sex: F(1, 164) = 233.13, p = 2.58e-33; age: F(2.06, 338.66) = 8209.73, p = 7.14e-290; condition x sex: F(1, 164) = 9.11, p = 0.0030; condition x age: (F(2.06, 338.66) = 12.70, p = 3.62e-06; condition x sex x age: F(2.06, 338.66) = 3.35, p = 0.035). Mice body weight was assessed on PND 11 (t-test with Bonferroni adjustment, F: p = 7.36e-15; M: p = 4.00e-23), 30 (F: p = 0.00029; M: p = 6.47e-10), 60 (F: n.s., M: p = 5.42E-09) and 110 (F: p = 0.00053; M: p = 2.65e-05). (C, E, G) Violin plots show mean (line) and individual mouse (symbol) percent time in open/inner zone over 10 minutes in (C) light dark box (t-test, t = 2.97, df = 172.69, p = 0.0034; n = 90 control, 85 LBN), (E) open field (t = 2.47, df = 101.67, p = 0.015; n = 56 control, 48 LBN), and (G) elevated zero maze tasks (n.s., n = 34 control, 33 LBN). (D, F, H) Kernel density estimates of percent time in open/inner zones of (D) light dark box, (F) open field, (H) and elevated zero maze tasks via Gaussian kernel. For all panels, *p <0.05, **p < 0.01, ****p < 0.0001. Mice are color-coded by rearing condition (control: gray, LBN: pink). Cartoons in panels (A, C, E, G) are made from modified BioRender templates (license Anderson, L. (2025) https://BioRender.com/wzzh1vz).

### LBN-reared mice show greater risk avoidance

Upon reaching early adulthood (PND ∼60-80), risk avoidance was assessed in LBN-reared and control mice using three well-established behavioral tasks: light-dark box, open field, and elevated zero maze. For all, the rodents’ innate drive to explore a novel environment was put into conflict with the drive for safety via the avoidance of exposed, bright zones [77]. Less time in open/inner zones was interpreted as greater risk avoidance. In both the LDB and OF tests, LBN-exposed mice spent less time in the open/inner zones than controls (Figure 1C, E). Density plots confirm a leftward shift percent time in open/inner scores among LBN-exposed mice (Figure 1D, F), In the EZM task, however, LBN-reared and control mice performed similarly (Figure 1G-H).

Though sex did not have a significant effect on performance in the LDB or OF tests (Figure S1A-B), a significant interaction between sex and condition emerged in the EZM task (Figure S1C). Interestingly, we found no significant correlations between time spent in the open zone of the LDB apparatus and time in the open/inner zones of either the OF or EZM apparatus (Figure S1D-E), suggesting that each task measures independent aspects of the risk-avoidance behavior.

### Increased accumbal dopamine D1-receptor binding in LBN-reared mice skews D1/D2-receptor ratio before alcohol exposure

To determine the effects of ELA on striatal dopamine receptor binding, D1– and D2-like receptor binding was compared between control and LBN-reared mice via autoradiography experiments (Figure 2A). No differences emerged in D2-like receptor binding, as quantified by the percent specific binding of [3H]Raclopride relative to control-reared mice (Figure 2B-C). However, LBN-reared mice showed significantly greater D1-like receptor binding in the NAc (Figure 2C) and a trend towards the same in the DMS (t-test, p-value = 0.099) compared to controls (Figure 2B). This shifted D1-to D2-like receptor ratios to trend higher among LBN-reared mice in the NAc (t-test, p-value = 0.087, Figure 2D).

**Figure 2.**
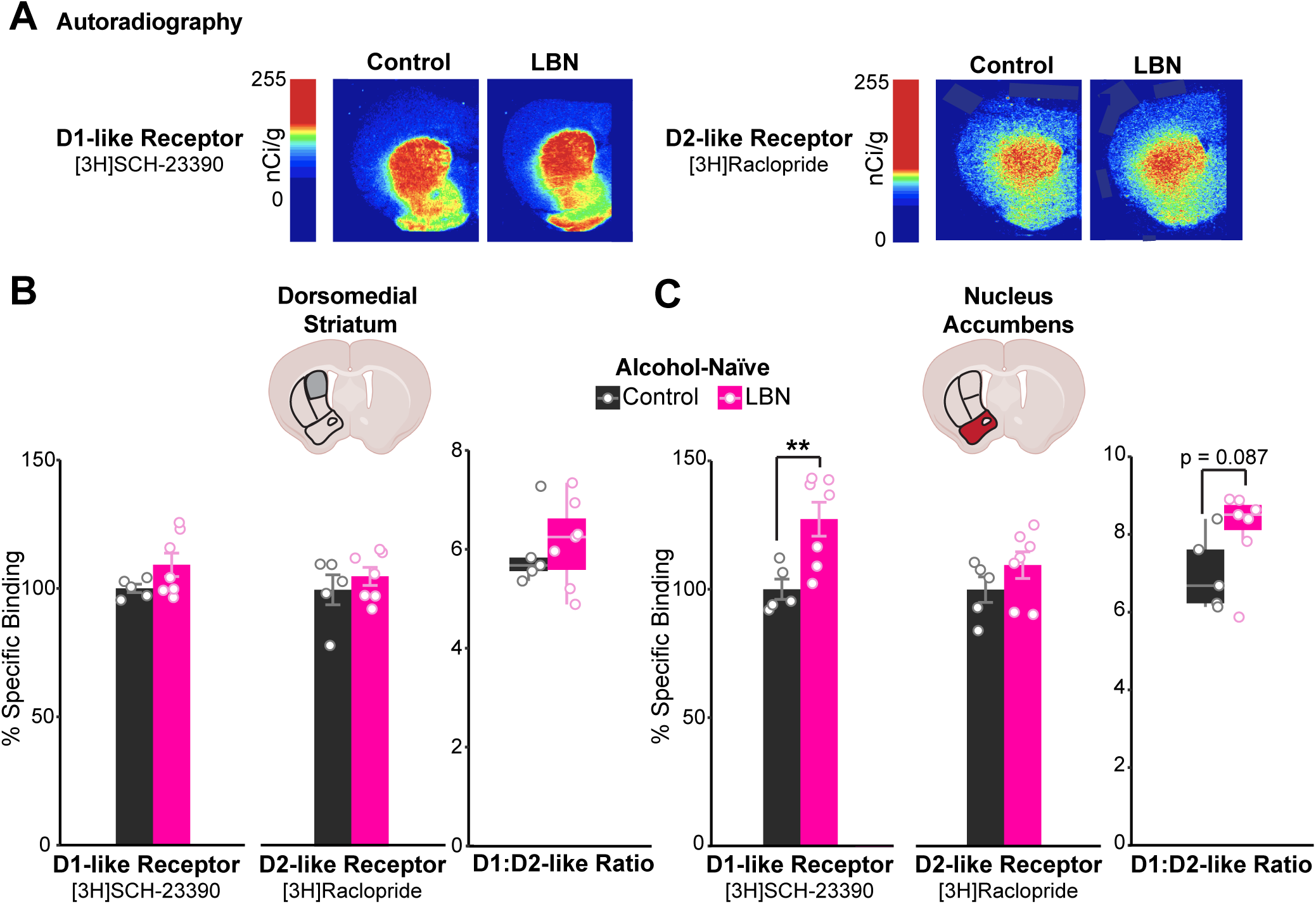
Higher striatal D1-like receptor binding among alcohol-naïve LBN-reared mice. **(A)** Representative images of [3H]SCH-23390 (left; D1-like receptor radioligand) and [3H]Raclopride (right; D2-like receptor radioligand) binding in coronal brain slices from alcohol-naïve control (left; n = 5) and LBN-reared (right; n = 7) mice. **(B, C)** Percent specific binding of [3H]SCH-23390 (left), [3H]Raclopride (middle), and their ratio (right) in the **(B)** DMS ([3H]SCH-23390: t-test, n.s.; [3H]Raclopride: n.s.; ratio: n.s.) and **(C)** NAc ([3H]SCH-23390: t-test, t = –3.54, df = 9.28, p-value = 0.0060; [3H]Raclopride: n.s., ratio: n.s.). Cartoons in panels **(B, C)** are made from modified BioRender templates (license Anderson, L. (2025) https://BioRender.com/wzzh1vz). **p < 0.01. Mice are color-coded by rearing condition (control: black, LBN: pink).

### LBN-exposed mice, especially males, drink more alcohol than controls in social-housed, voluntary operant drinking task

We hypothesized that preexisting alterations to dopamine D1-receptor availability in the NAc may increase vulnerability to AUD-like behaviors in LBN-reared mice. Operant, intermittent alcohol drinking was assessed during social housing using the IntelliCage System in which up to 16 same-sex mice/cage received access to 20% alcohol every other day on an FR-3 schedule (Figure 3A). Due to differences in body weight (Figure 1B), alcohol consumption is presented as licks per gram of body weight.

**Figure 3.**
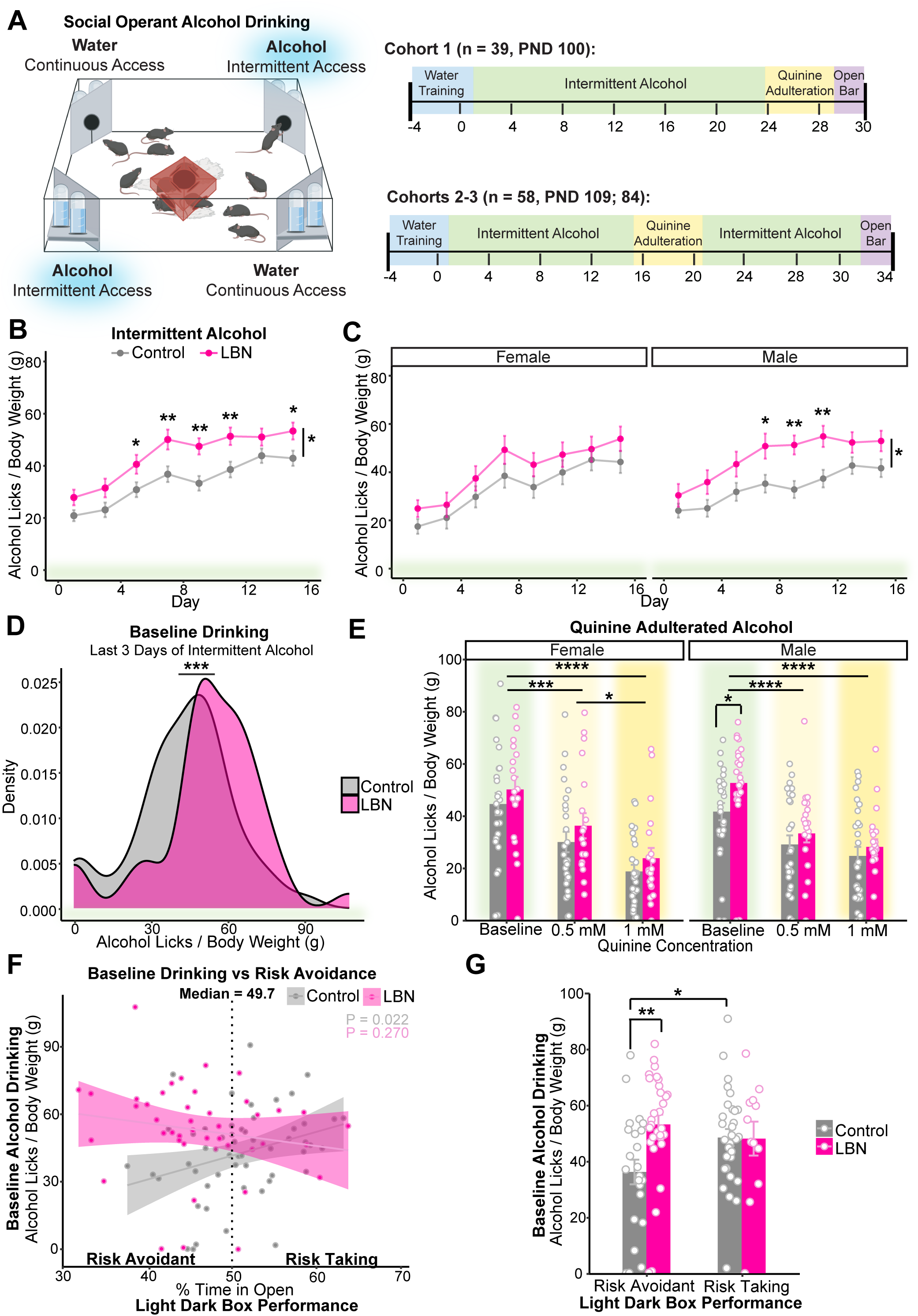
LBN-reared mice drink more alcohol than controls, especially males and those that are more risk avoidant. **(A)** Schematic and experimental timeline. Cartoon made from modified BioRender templates (license Anderson, L. (2025) https://BioRender.com/wzzh1vz). **(B, C)** Mean alcohol licks per day normalized by body weight (g; measured weekly) during 8 days of intermittent alcohol access **(B)** collapsed by sex (Mixed ANOVA, condition: F(1, 95) = 8.58, p = 0.0040; day: F(3.86, 367.13) = 43.86, p = 2.63e-29; t-test with Bonferroni adjustment, day 5: p = 0.035; 7: p = 0.0069; 9: p = 0.0011; 11: p = 0.0059; 15: p = 0.019; n = 54 control, 43 LBN) and **(C)** separated by sex (F: left; Mixed ANOVA, day: F(3.22, 141.51) = 21.61, p = 4.11e-12; n = 26 control, 20 LBN; M: right; Mixed ANOVA, condition: F(1, 49) = 6.69, p = 0.013; day: F(3.94, 193.24) = 23.02, p = 2.02e-15; t-test with Bonferroni adjustment, day 7: p = 0.016; 9: p = 0.0011; 11: p = 0.0034; n = 28 control, 23 LBN). Points show mean and bars show SEM. **(D)** Kernel density estimation via Gaussian kernel of pre-adulteration baseline drinking in the last three alcohol days of intermittent drinking before quinine adulteration (ANOVA, condition: F(1) = 11.65, p = 0.00079; n = 54 control, 43 LBN). **(E)** Mean alcohol licks per day normalized by body weight (g) among female (left) and male (right) mice before and during quinine adulteration (0.5 mM, 1 mM) (Mixed ANOVA, day: F(1.86, 172.63) = 155.35, p = 1.08e-37; day x sex: F(1.86, 172.63) = 4.13, p = 0.020; F: t-test with Bonferroni adjustment, 0.5 mM vs baseline: p = 0.00084; 1 mM vs baseline: p = 2.99E-09; 0.5 mM vs 1 mM: p = 0.016; n = 26 control, 20 LBN; M: t-test with Bonferroni adjustment, baseline: p = 0.037; 0.5 mM vs baseline: p = 4.48E-05; 1 mM vs baseline: p =9.53E-08; n = 28 control, 23 LBN). Bars showing mean and SEM are overlaid with symbols showing data from individual mice. **(F)** Correlation between percent time in open during the light dark box test and baseline alcohol drinking among control (linear regression, y = –11.4 +106x, *R*^2^ = 0.08, F(1, 51) = 5.58, p = 0.022; n = 53) and LBN-reared (y = 76.2 – 50.6x, *R*^2^ < 0.01, F(1, 41) = 1.25, p = 0.27; n = 43) mice. Symbols represent individual data points and shading shows 95% confidence interval. Dashed line shows median value used to split population into more risk-avoidant (% time in open < 49.7) and more risk-taking (% time in open > 49.7) subgroups. **(G)** Alcohol licks per body weight (g) during pre-adulteration baseline among more risk-avoidant and more risk-taking subgroups (ANOVA, condition: F(1) = 6.84, p = 0.010; condition x median split: F(1) = 4.81, p = 0.031; Tukey’s, risk-avoidant: p = 0.0052; risk-avoidant: n = 23 control, 31 LBN; risk-taking: n = 31 control, 12 LBN). Bar showing mean and SEM are overlaid with symbols showing data from individual mice. For all panels, *p <0.05, **p < 0.01, ***p<0.001, ****p < 0.0001. Mice are color-coded by rearing condition (control: gray, LBN: pink).

During intermittent, operant access, LBN-reared mice drank significantly more alcohol than controls (Figure 3B). Though significant interactions with sex did not emerge, given the well-described sex difference in alcohol consumption in mice [78,79] including LBN-reared mice [42–44], we also analyzed drinking separately by sex. Notably, we found that rearing condition only emerged as a significant main effect among male mice (Figure 3C).

Interestingly, during alcohol access, LBN-reared mice also drank more water than controls (Figure S2A). This effect was primarily driven by females (Figure S2B). This contrasts with data from alcohol-naïve mice (Figure S2C) among which differences in water consumption were more pronounced among males (Figure S2D). Though the source of this polydipsia is unclear, it may be connected to differences in baseline risk avoidance and stress exposure [80–82].

To assess whether differences in drinking behavior were unique to an FR-3 schedule, cohorts one, two and three underwent a single day of FR-0 alcohol and water drinking. Consistent with our FR-3 results, during the “Open Bar” module, LBN-reared mice trended towards higher daily alcohol consumption than controls (Figure S2E).

### LBN-reared and control mice suppressed drinking similarly in response to quinine adulteration

We predicted that LBN-reared mice may show more punishment-insensitive drinking. To assess this, 20% alcohol was adulterated with quinine (0.5 mM, 1 mM), a bitter-tasting substance, on consecutive alcohol-drinking days. Pre-adulteration baseline drinking was quantified for each mouse by averaging daily alcohol licks per gram of body weight for three days before adulteration (Figure 3D).

Contrary to our hypothesis, LBN-exposed and control mice suppressed their alcohol consumption similarly in response to quinine adulteration (Figure 3E). A dose x sex interaction emerged such that only females showed a significant reduction between 0.5– and 1-mM concentrations. Similar results emerged when we calculated percent reduction in baseline drinking during quinine adulteration (Figure S2F).

Cohorts two and three were returned to alcohol drinking following adulteration. Female mice rebounded their drinking faster than males after adulteration, but no differences emerged between control and LBN-reared mice (Figure S2G).

As a control, quinine-adulterated water consumption was measured among alcohol-naïve mice. We found equal sensitivity to quinine adulteration between control and LBN-reared mice (Figure S2H).

### Differences in alcohol consumption between control and LBN-reared mice driven by more risk-avoidant mice

We hypothesized that increases in risk avoidance among LBN-reared mice (Figure 1C-F) may underlie increases in social, operant alcohol drinking. To investigate this, we correlated percent time in the open section of the light-dark box with pre-adulteration alcohol drinking (Figure 3F). A median-split (median = 49.7%) revealed that differences in baseline drinking between LBN-reared and control mice were disproportionately driven by mice with greater risk avoidance (Figure 3G).

### LBN-reared and control mice have similar sensitivity to the anxiolytic potency of alcohol

We hypothesized that differences in sensitivity to the anxiolytic effects of alcohol may promote alcohol drinking among LBN-reared mice with greater baseline risk avoidance. On PND ∼65-75, mice underwent repeated testing in an elevated zero maze, once after injection of saline and once after alcohol (1.2 g/kg, i.p.; order counterbalanced), an approach that our group recently found to be effective in measuring individual differences in alcohol relief [64]. The stability of EZM performance run one week apart was found by our group and others to be satisfactorily consistent (83,64). We also attempted to measure anxiolytic potency of alcohol in a repeated open field task but did not see an anxiolytic effect (1.2 g/kg or 1.7 g/kg, i.p.; Figure S3G).

Ultimately, control and LBN-reared mice performed similarly on the repeated EZM task (Figure S3A). Sensitivity to the anxiolytic effects of alcohol (quantified as Alcohol Effect = % Time in Open_Alcohol_-% Time in Open_Saline_) was strongly correlated with baseline risk avoidance (quantified as percent time in the open after saline; Figure S3B). However, even within more risk avoidant (“Below Median”) and risk-taking (“Above Median”) subgroups, alcohol effect was similar between LBN-reared and control mice (Figure S3C). LBN-reared were roughly equally represented in below and above median groups (Figure S3D).

When we compared sensitivity to the anxiolytic effects of alcohol to baseline alcohol drinking in the Intellicages, similar trends emerged among control and LBN-reared mice (Figure S3C), however females with greater sensitivity to the anxiolytic effects of alcohol had a higher alcohol consumption in the intermittent alcohol task (Figure S3B).

### Greater alcohol stimulation in LBN-reared females, while stronger sedation in LBN-reared males

Given that striatal D1-receptor activation is known to play a key role in the acute stimulatory effects of alcohol [52,55,64], we hypothesized that enhanced stimulation may be another mechanism driving alcohol drinking among LBN-reared mice. Ethanol-induced locomotion was assessed via repeated testing in an open field box following subsequent saline and alcohol injection (1.2 or 1.7 g/kg, i.p.) across two days, with injection order counterbalanced (Figure 4A). Distance travelled (m) per 5-minute time bin was quantified via automated video tracking. Injection-induced locomotion was calculated as follows: Δ Distance Travelled = Post-Inj_5 min_ − Post-Inj_Avg 15 min_. Differences in distance travelled > 0 were interpreted as stimulatory while those < 0 were interpreted as sedatory. Pre-injection locomotion was found to be similar among LBN-reared and control mice (Figure S3H). Due to sex differences in response to injection (Mixed ANOVA, sex: F(1, 192) = 6.10, p = 0.014; sex x condition: F(1, 192) = 10.70, p = 0.0010; sex x injection: F(2, 192) = 6.35, p = 0.0020), we have chosen to analyze the data from male and female mice separately in this task.

**Figure 4.**
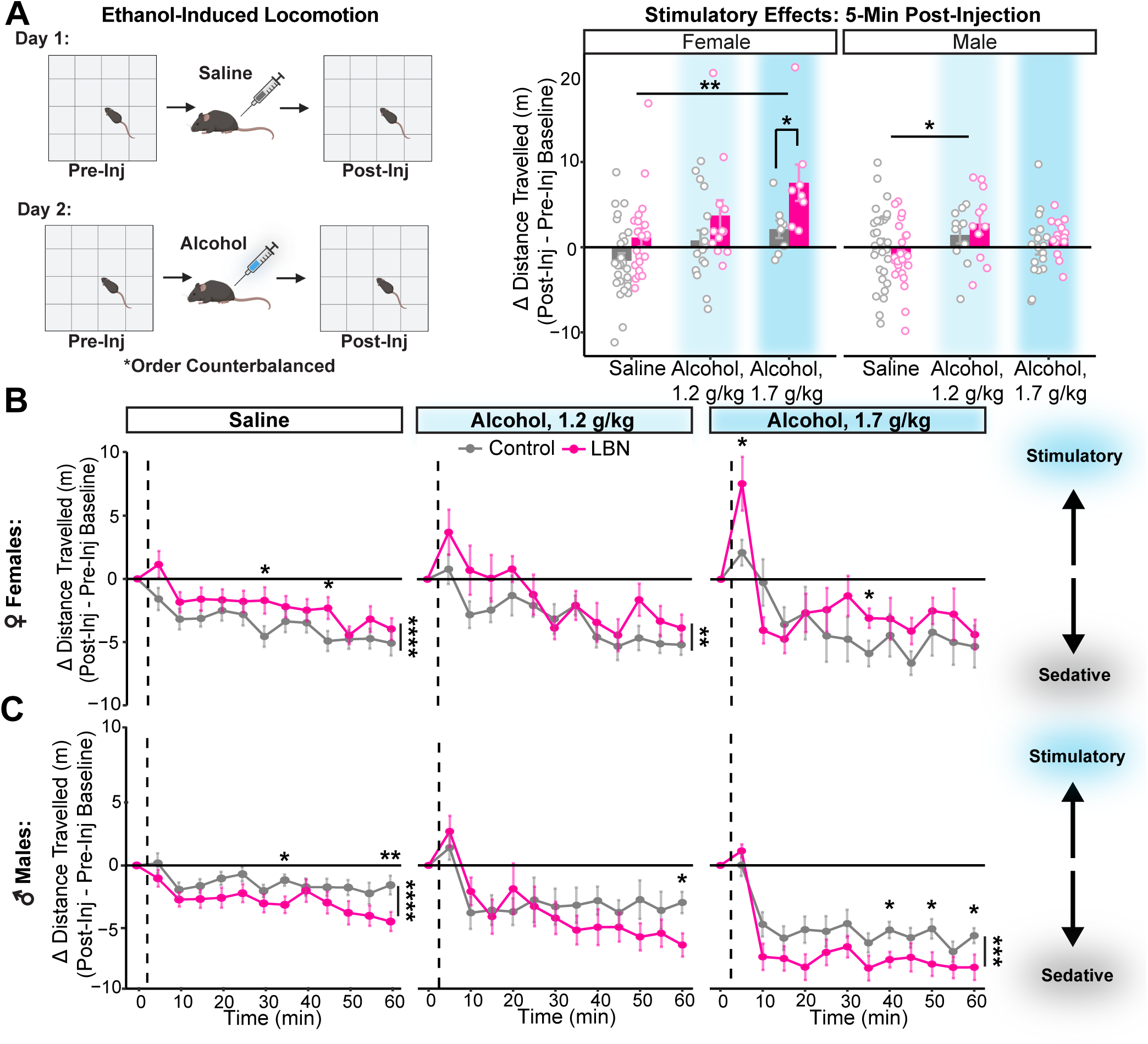
LBN-reared females show increased sensitivity to the stimulatory effects of alcohol. **(A)** Experimental design. Cartoon made from modified BioRender templates (license Anderson, L. (2025) https://BioRender.com/wzzh1vz). Distance travelled (m) five minutes after injection normalized to pre-injection baseline among female (left) and male (right) mice (F: ANOVA, condition: F(1) = 10.32, p = 0.0018; injection: F(2) = 6.69, p = 0.0020; t-test with Bonferroni adjustment, saline vs alcohol, 1.7 g/kg: p = 0.0022; alcohol, 1.7 g/kg: p = 0.037; n = 26 control, 21 LBN; M: ANOVA, injection: F(2) = 3.50, p = 0.034; t-test with Bonferroni adjustment, saline vs alcohol, 1.2 g/kg: p = 0.0265; n = 30 control, 26 LBN). Bars show mean and SEM are overlaid with data from individual mice. **(B, C)** Points showing mean (and SEM) distance travelled per 5-minute time bin normalized by pre-injection baseline for 60 minutes after injection with saline or alcohol (1.2 g/kg or 1.7 g/kg) among **(A)** female (mixed ANOVA, condition: F(1, 87) = 5.09, p = 0.027; time bin: F(9.03, 785.94) = 26.11, p = 1.08e-39; injection x time bin: F(18.07, 785.94) = 2.60, p = 0.00030; t-test with Bonferroni adjustment, saline: p = 4.75E-05, saline(bin 6) = 0.036, saline(bin 9) = 0.0334; alcohol, 1.2 g/kg: p = 0.0020; alcohol, 1.7 g/kg(bin 7) = 0.046; saline: n = 26 control, 21 LBN; alcohol, 1.2 g/kg: n = 18 control, 12 LBN; alcohol, 1.7 g/kg: 8 control, 8 LBN) and **(B)** male mice (mixed ANOVA, condition: F(1, 105) = 5.67, p = 0.019; injection: F(2, 105) = 17.98, p = 1.92e-07, time bin: F(8.83, 926.7) = 41.60, p = 1.18e-61, condition x time bin: F(8.83, 926.7) = 2.83, p = 0.00030, injection x time bin: F(17.65) = 5.41, p = 5.38e-12; t-test with Bonferroni adjustment, saline: p = 1.20E-06, saline(bin 7): p = 0.017, saline (bin 12): p = 0.0098; alcohol, 1.2 g/kg(bin 12): p = 0.015; alcohol, 1.7 g/kg: p = 0.00018, alcohol, 1.7 g/kg(bin 8): p = 0.016, alcohol, 1.7 g/kg(bin 10): p = 0.029, alcohol, 1.7 g/kg(bin 12): p = 0.034; saline: n = 30 control, 26 LBN; alcohol, 1.2 g/kg: n = 11 control, 11 LBN; alcohol, 1.7 g/kg: 18 control, 15 LBN). For all panels, *p <0.05, **p < 0.01, ***p<0.001, ****p < 0.0001. Mice are color-coded by rearing condition (control: gray, LBN: pink).

Among female mice, LBN-exposed mice showed greater post-injection locomotion than controls (Figure 4B). Direct comparison of the acute stimulatory effects observed 5 minutes post-injection revealed that though both control and LBN-reared females were sensitive to the stimulatory effects of alcohol, LBN-reared females showed greater stimulation than controls (Figure 4A). Further, LBN-exposed females trended towards having less sensitivity to the sedative effects of alcohol, 45-60 minutes post-injection (ANOVA, condition: F(1) = 3.86, p = 0.053; Figure S3I).

In contrast, LBN-exposed males showed lower post-injection locomotion than controls (Figure 4C). Male LBN-reared mice reacted similarly to the stimulatory effects of alcohol (Figure 4A) but showed greater sedation than controls following all injections, including saline (Figure S3I).

### Alcohol drinking lowers striatal D1– and D2-like receptor binding, ultimately normalizing D1/D2-like receptor ratios in LBN-reared and control mice

To determine how alcohol exposure interacts with ELA to modulate dopamine D1– and D2-receptor balance throughout the striatum, we assessed D1– and D2-like receptor binding via autoradiography in age-matched LBN-reared and control mice following alcohol exposure (2x alcohol injection + voluntary, operant intermittent alcohol drinking). Results from alcohol-exposed mice were normalized to region-specific averages among alcohol-naïve controls and are shown alongside the results from Figure 2 (Figure 5A-F).

**Figure 5.**
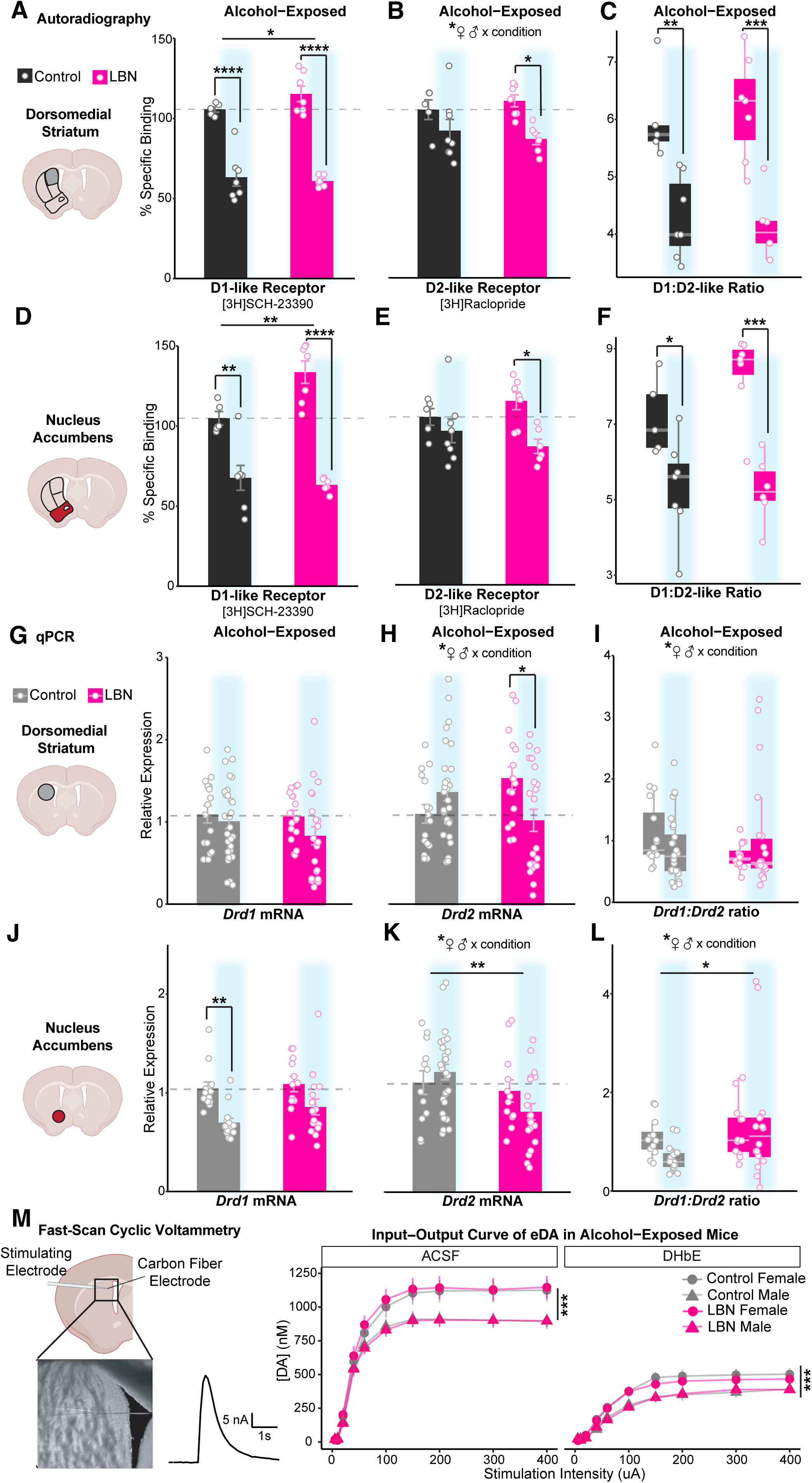
Alcohol exposure interacts with adversity and sex to lower striatal dopamine D1 and D2 receptor transcription and translation, especially in LBN-reared males, ultimately resulting in similar D1/D2 receptor ratio. Percent specific binding of **(A, D)** [3H]SCH-23390 (left), **(B, E)** [3H]Raclopride (middle), and their **(C, F)** ratio (right) among alcohol-naïve and – exposed (blue highlight) control (black; n = 5 naïve, 7 exposed) and LBN-reared (pink; n = 7 naïve, 7 exposed) mice in the **(A, B, C)** dorsomedial striatum ([3H]SCH-23390: ANOVA, alcohol: F(1) = 127.08, p = 2.28e-10; condition: F(1) = 4.68, p = 0.042; Tukey’s, control: p = 6.20E-06; LBN: p = 5.35E-08; [3H]Raclopride: ANOVA, alcohol: F(1) = 13.39, p = 0.0018; condition x sex: F(1) = 5.42, p = 0.032; Tukey’s, LBN: p = 0.018; ratio: ANOVA, alcohol: F(1) = 39.08, p = 3.36e-06; Tukey’s, control: p = 0.0042; LBN: p = 0.00040) and the **(D, E, F)** nucleus accumbens ([3H]SCH-23390: ANOVA, alcohol: F(1) = 77.21, p = 1.77e-08; condition: F(1) = 8.50, p = 0.0083; condition x alcohol: F(1) = 7.08, p = 0.015; Tukey’s, control: p = 0.0025; LBN: p = 3.14E-07; [3H]Raclopride: ANOVA, alcohol: F(1) = 8.90, p = 0.0069; Tukey’s, LBN: p = 0.019; ratio, ANOVA: alcohol: F(1) = 31.98, p = 1.3e-05; Tukey’s, control: p = 0.040; LBN: 0.00029). Relative expression of (G, J) *Drd1*, (H, K) *Drd2* and their **(I, L)** ratio among alcohol-naïve and –exposed (blue highlight) control (gray; n = 16 naïve, 30 exposed) and LBN-reared (pink; n = 16 naïve, 26 exposed) mice in the **(G, H, I)** dorsomedial striatum (*Drd1*: ANOVA, n.s.; *Drd2*: ANOVA, condition x alcohol: F(1) = 10.71, p = 0.0016; condition x sex: F(1) = 10.68, p = 0.0016; Tukey’s, LBN: p = 0.0121; ratio: ANOVA, condition x alcohol: F(1) = 3.99, p = 0.049; condition x sex: F(1) = 7.90, p = 0.0063; Tukey’s, n.s.) and **(J, K, L)** nucleus accumbens (*Drd1*: ANOVA, alcohol: F(1) = 16.71, p = 0.00016; Tukey’s, control: p = 0.0066; *Drd2*: ANOVA, condition: F(1), 10.51, p = 0.0019; condition x sex: F(1) = 5.90, p = 0.018, alcohol x sex: F(1) = 4.67, p = 0.035; ratio: ANOVA, condition: F(1) = 5.02, p = 0.030; condition x sex: F(1) = 5.30, p = 0.026). For panels **(A-L)**, mean (bars) and SEM (error bars) are overlaid with data from individual mice. **(M)** Electrically-evoked dopamine signals in the DMS of control (gray; n = 66 slices/13 animals) and LBN-reared (pink; 61 slices/12 animals) mice in ACSF and after bath application of 1 μM DHßE (Mixed ANOVA, drug: F(1, 201) = 234.03, p = 1.52e-35; sex: F(1, 201) = 14.70, p = 0.00017; sex x stimulation: F(1.88, 377.07) = 12.65, p = 8.26e-06). Points show mean value and bars show SEM. For all panels, *p <0.05, **p < 0.01, ***p<0.001, ****p < 0.0001. Cartoons in panels **(A, D, G, J, M)** are made from modified BioRender templates (license Anderson, L. (2025) https://BioRender.com/wzzh1vz).

In both the NAc and DMS, alcohol exposure interacted with condition to strongly decrease specific binding of D1-like receptors, especially among LBN-reared mice (Figure 5A, D). This robust decrease negated previous differences between LBN-reared and control mice, particularly in the NAc.

Alcohol exposure also reduced binding of D2-like receptors, though to a slightly lesser extent. Once again, this effect was most pronounced in LBN-reared mice (Figure 5B, E). Interestingly, a significant interaction between sex and condition emerged in the DMS. Though our sample size is too small to make definitive claims about sex differences, preliminary trends suggest that LBN-reared males may experience the largest reduction in D2-like receptor density following alcohol drinking (Figure S4A).

Alcohol-induced changes to transcription and translation resulted in large reductions to D1-to D2-like receptor ratio. Driven primarily by reductions in D1-like receptor binding, which preferentially impacted LBN-reared mice, alcohol exposure negated pre-existing differences in receptor ratio between LBN-reared and control mice (Figure 5C, F). Interestingly, D1-to D2-like receptor ratio trended towards a positive correlation with baseline alcohol drinking among LBN-reared mice (Figure S4F-G).

### Alcohol affects *Drd1* and *Drd2* transcription differently in LBN-reared and control mice, yet transcriptional changes do not fully account for changes in receptor density

Having found robust changes to striatal D1– and D2-like receptor binding after alcohol exposure, we sought to determine the extent to which alcohol’s effects were transcriptionally regulated. Using bulk mRNA quantification of DMS and NAc tissue punches, we assessed changes in *Drd1* and *Drd2* transcription after alcohol drinking among control and LBN-reared mice relative to alcohol-naïve controls (Figure 5G-L).

Quantitative polymerase chain reaction (qPCR) analysis of *Drd1* expression found modest reductions in expression that do not fully account for the large reductions seen post-transcriptionally. In the DMS, though Drd1 trended lower, expression was statistically comparable (Figure 5A). In the NAc, alcohol-exposed mice had lower relative expression of *Drd1* than naïve mice, though no differences emerged between rearing conditions (Figure 5C). Alcohol exposure also affected expression of *Drd2* (Figure 5B, D). Alcohol interacted with rearing condition and sex to produce significantly lower *Drd2* expression in LBN-reared mice, especially males, in both the DMS (Figure 5B, S4B) and NAc (Figure 5D, S4C). When analyzed separately by sex, only LBN-reared males showed a significant reduction in *Drd2* expression (Figure S4B-C).

In the DMS, alcohol exposure negated preexisting differences in *Drd1:Drd2* ratio between LBN-reared and control mice, leading to roughly comparable values among mice (Figure 5I). However, inconsistent with autoradiography findings, LBN-reared mice had higher *Drd1:Drd2* ratios than controls in the NAc, especially among males (Figure 5L, 5D-E). This was driven by the drop in *Drd2* expression following alcohol exposure, which mirrors that seen post-transcriptionally among LBN-reared mice.

### Sex differences in striatal dopamine release capacity after alcohol drinking are independent of rearing condition

To confirm the functional similarity of striatal dopamine signaling in alcohol-exposed LBN-reared and control mice, we assessed dopamine release capacity in the DMS of *ex vivo* brain slices via FSCV (Figure 5M). The amplitude of the electrically-evoked dopamine signals were assessed at baseline in ACSF, as well as after the bath application of a nicotinic receptor blocker, DHßE, to determine if cholinergic interneuron contribution was comparable between control and LBN-reared mice. Surprisingly, sex but not condition had a main effect on electrically evoked dopamine, such that females had higher electrically-evoked dopamine than males (Figure 5M). Response to DHßE was similar between groups and sexes.

## DISCUSSION

In this study we sought to understand the neurobiological alterations induced by a brief yet potent early-life perturbation, and the lasting effects on risk and social alcohol drinking. Mice exposed to ELA via the limited bedding and nesting paradigm had greater baseline risk avoidance and voluntarily consumed more alcohol than their control counterparts. In line with previous reports, differences in alcohol consumption were especially pronounced among LBN-reared males [42,44]. Further, we found that LBN-rearing increases dopamine D1-receptor binding in the NAc, ultimately skewing the ratio of D1-to D2-receptors higher. However, after voluntary alcohol exposure, robust alcohol-induced effects on D1– and D2-receptor binding, led to comparable receptor densities in control and LBN-reared mice. We hypothesize that these alcohol-induced changes in receptor density occur via both transcriptional and post-transcriptional mechanisms, as they are only partially matched by the expression patterns. In all, our results uncover specific neurobiological mechanisms that promote alcohol consumption after LBN-rearing.

### Effects of LBN-rearing on striatal reward circuitry and related behavioral outcomes

Broadly, rodent models of early life adversity are known to affect the brain via neuroinflammatory and epigenetic mechanisms following increases in hypothalamic-pituitary axis and sympathetic nervous system activity [84–87]. In humans, these ELA-induced changes to neural circuitry affect later reward processing [5–11], ultimately conferring vulnerability to both mood disorders and SUDs [14,15,8,20,19,21]. Preclinical models are ideal for disaggregating genetic and environmental influences on development because of access to isogenic animals and tighter control over environmental conditions. The extent to which models of ELA in rodents mirror observations in clinical populations varies by paradigm and investigation [39,27,87]. The LBN paradigm alters the quality of maternal care, inducing more fragmented and unpredictable caregiving [71–75,88,76,89]. The unpredictability of sensory cues is thought to be a key component of the adversity induced by this paradigm [11,73], enhancing its translatability [90]. In our hands, we found profound developmental consequences of LBN-rearing (PND 3-10) among genetically identical, C57BL/6J mice. Consistent with previous reports, pups exposed to LBN-rearing weighed significantly less than controls into adulthood (Figure 1B) [79,82,95].

As hypothesized, LBN-rearing robustly altered the binding of striatal dopamine receptors. LBN-reared mice exhibited higher specific binding of D1-like receptors (Figure 2). This was especially true in the NAc, where elevated D1-like receptor binding pushed D1-to D2-like receptor binding ratios to trend higher among LBN-reared mice. Though previous studies of ELA models have documented transcriptional changes to the brain’s reward circuitry [46,47], including a reduction of D1-receptor expression in the NAc [45], ours is unique in assessing D1– and D2-like receptors post-transcriptionally using autoradiography. We are also the first to study the effects of ELA in the DMS, a region involved in cognitive flexibility and impaired in AUD [48,49]. Transcriptionally, we found comparable *Drd1* expression among alcohol-naive mice, suggesting that increases in D1-like receptor binding among LBN-reared mice may arise post-transcriptionally (Figure 5G-L). More study is needed to clarify the specific post-transcriptional mechanisms by which ELA acts.

Striatal D1-receptor activation plays a key role in the acute stimulatory effects of alcohol [52,55,64], and D1-receptor-expressing striatal neurons promote AUD-like behaviors, more broadly [50–55,64]. This is in line with clinical reports linking higher stimulation and lower sedation in response to alcohol with increased AUD vulnerability [56–63]. Further, we previously found that mice with altered D1/D2-receptor ratios exhibit greater risk avoidance and heightened alcohol relief [64], consistent with pre-clinical [92] and clinical reports connecting low D2-receptor binding with enhanced SUD vulnerability [56–63]. Given our findings that LBN-reared mice had greater binding of striatal D1-like receptors and skewed D1/D2-receptor ratios, we predicted that enhanced sensitivity to alcohol’s acute stimulatory and anxiolytic effects may be driving higher alcohol consumption among LBN-reared mice.

We were the first to use social housing in the study of alcohol drinking among ELA-exposed pups, an important advancement in reducing the confounding effect of a second-adverse event induced by social isolation. Confirming findings among individually-housed mice [44], we found that LBN-reared mice voluntarily consumed more alcohol than controls in a social drinking task (Figure 3B-D). This effect was especially pronounced among LBN-exposed males, corroborating previous reports [42] and clinical trends [8,14,15,19–21,25,93]. LBN-reared mice did not engage in more punishment-insensitive drinking than controls under the conditions tested (Figure 3E).

Though ELA is robustly associated with the development of anxiety disorders in the clinical population [16,17,94,95], mixed findings have emerged in studies of rodents exposed to the LBN paradigm [27,75,88,89]. In our study, LBN-reared mice showed increased avoidance of open/inner zones in two of three risk avoidance tasks, at very robust sample sizes (LDB: n = 173, OF: n = 102; Figure 1C-F). Clinically, ELA followed by the onset of depression, anxiety, or other psychological disorders increases the risk for comorbid binge drinking and AUD in adulthood [15,37]. In line with this and our proposed role of striatal dopamine receptor balance, we found that differences in alcohol drinking were more pronounced in LBN-reared mice that were more risk avoidant in the light-dark box (Figure 3F-G). However, when we directly measured alcohol relief, LBN-exposed mice showed similar sensitivity to alcohol’s anxiolytic potency (Figure 4A-D). In our ethanol-induced locomotion task, only female LBN-reared mice showed heightened locomotion following alcohol (Figure 4E-F), in line with Morningstar [44]. Though LBN-reared males trended towards greater sedation than controls after alcohol (1.7 g/kg), Morningstar found less sedation among LBN-reared males at a higher dose of alcohol (4 g/kg) [44]. Our findings of greater sensitivity to the stimulatory effects of alcohol among females are consistent with studies linking D1-receptor activation to the stimulatory effects of alcohol in the NAc [52] and DMS [55,64] and support our hypothesis that greater striatal D1-receptor activation in LBN-reared mice drives increases to alcohol drinking.

### Alcohol interactions with adversity-mediated changes to striatal dopamine receptors

Though others have found effects of adulthood experience on adversity-induced changes to transcription [11,28,32], to our knowledge, the impacts of alcohol were yet uninvestigated. We found robust effects of alcohol exposure (2x alcohol injection + voluntary alcohol drinking) on dopamine receptor binding across the striatum. Alcohol lowered D1-receptor binding by over 40% across the striatum via a post-transcriptional mechanism, independently of the rearing history (Figure 5A, D). These findings, while novel, are consistent with general reports on the effects of alcohol exposure on striatal dopamine receptors [96–99]. Proportionally, alcohol suppression was larger in LBN-reared mice such that after alcohol drinking, D1-like receptor binding was comparable between control and LBN-reared mice. Though *Drd1* mRNA levels also decreased after alcohol (Figure 5G, J), changes in expression do not fully account for our autoradiography results. We speculate that mechanisms downstream of gene expression are also at work.

Alcohol also lowered D2-like receptor binding, though only by ∼15% and mainly in LBN-reared mice (Figure 5B, E). Alcohol suppression of D2-like receptor binding was accompanied by lower *Drd2* mRNA levels selectively in LBN-reared males (Figure 5H, K; Figure S4B-C). Future study is needed to clarify the role of sex-related factors on D2-like receptor transcription and translation after adversity.

Ultimately, through a combination of transcriptional and post-transcriptional regulation, alcohol lowered striatal dopamine receptors levels to negate preexisting differences in D1-to D2-like receptor ratios among LBN-reared mice. Functionally, dopamine release capacity was also similar between control and LBN-reared mice after alcohol exposure (Figure 5M). Due to the voluntary design of our alcohol drinking task and the fact the LBN reared mice consumed more alcohol, we cannot rule out alcohol x condition interactions resulting from increased alcohol consumption rather than the effects of ELA. DMS and NAc receptor ratios are positively correlated with alcohol drinking (Figure S4F-G), counter to the overall trend we find in response to alcohol. Further studies are needed to identify the gene repressor(s) at play and rule out latent effects of LBN-rearing.

Though we did not perform any behavioral studies to assess the outcomes of these alcohol-induced changes to striatal reward circuitry, results from Morningstar suggest that alcohol drinking may reverse the effect of ELA on anxiety-like behavior in males, which would be consistent with our autoradiography results [44].

In conclusion, our findings suggest that early life adversity alters striatal reward circuitry to increase D1-receptor binding, especially in the NAc. Though this upregulation may initially be adaptive, we hypothesize that increased striatal D1-like receptor binding enhances the sensitivity of LBN-reared mice to alcohol’s stimulatory effects to promote alcohol drinking, especially among mice with greater risk avoidance. In turn, alcohol has robust effects on dopamine receptor binding, lowering D1– and D2-receptor binding to effectively normalize ratios between control and LBN-reared mice after voluntary alcohol drinking. Our results support the interacting role of sex-related factors on the effects of ELA, consistent with previous literature [24–28]. However, future work is needed to parse out their underlying mechanisms. In all, the findings of this study point towards an interplay between genetics, sex-related factors, and experience in both early life and adulthood in the mediation of AUD vulnerability. Future studies should test whether these mechanisms translate to clinical populations and further explore the role of D1-receptor upregulation in the promotion of AUD-like behaviors following early life adversity.

## DATA AVAILABILITY STATEMENT

All data and code are accessible on Mendeley. Reserved DOI: 10.17632/5ygmcc6cm4.1.

## ACKNOWLEDGEMENTS

We would like to thank the NIMH IRP Rodent Behavioral Core for their help with behavioral testing and Drs. Miriam Bocarsly and Yan Leng for their technical advice. Additionally, we would like to thank Mikal Armstrong, our former HI-STEP 2.0 intern, for her help running Intellicage experiments. The contributions of the NIH authors were made as part of their official duties as NIH federal employees, are in compliance with agency policy requirements, and are considered Works of the United States Government. However, the findings and conclusions presented in this paper are those of the authors and do not necessarily reflect the views of the NIH or the U.S. Department of Health and Human Services.

## AUTHOR CONTRIBUTIONS

Conceptualized by L.G.A., V.A.A., R.B., and M.M. All experimentation except autoradiography conducted by L.G.A. Autoradiography conducted by A.T. Technical support by R.B. Formal analysis by L.G.A., A.T., and R.B. Writing and figure design by L.G.A. Review and editing by all authors. Funding acquisition by V.A.A. and M.M.

