## Supplement for "Early life adversity increases striatal dopamine D1 receptor density and promotes social alcohol drinking in mice, especially males"

### SUPPLEMENTARY MATERIALS AND METHODS

**Animals:** All experimental procedures were approved by the NIH Institutional Animal Care and Use Committee and complied with Public Health Service policy on the humane care and use of laboratory animals. Mice were classified as “male” or “female” based on external genitalia and/or anogenital distance. All mice were group-housed on a 12 h:12 h light cycle (6:30 on, 18:30 off) with dry-bulb temperature of 20-26 °C and relative humidity in the range of 30-70%. Mice had *ad libitum* access to standard rodent chow and water.

**Limited Bedding and Nesting Paradigm:** After arrival, timed pregnant dams (gestational day 17, n = 33; C57BL6/J background, JAX: 000664) were rehoused into standard housing conditions and checked twice daily for births. Among all cohorts (5 cohorts, 6-7 dams/cohort), timed-pregnant dams gave birth within 24 hours of each other. On PND 3, pups were removed from their home cages and organized by sex in euthermic holding cages for cross-fostering. Dams were randomly assigned to LBN or control conditions and were rehoused in new cages. Pups were rubbed with bedding material from their new dam (to reduce the risk of cannibalization) and then randomly assigned to control or LBN cages, aiming for equivalent litter sizes (usually 6-7 mice) and 1:1 sex ratios. LBN cages had limited corn cob bedding and were equipped with a fine-gauge, plastic-coated aluminum mesh platform (McNichols Co., 4700313244, Livermore, CA, USA) set approximately 2.5 cm above the cage floor with half of a single cotton Nestlet square (Ancare, NES3600, Bellmore, NY, USA). Control cages contained standard amounts of corn cob bedding and one full cotton Nestlet square. Both control and LBN cages were of the same standard size, 31 (l) × 13 (w) × 14 (h) cm. Control and LBN cages remained undisturbed for 7 days (PND 3-10) during which they were checked at least once daily. Any pups found failing to thrive (as indicated by lack of milk spot, dehydration, neglect, etc.) were euthanized. On PND 11, both control and LBN dams and their pups were rehoused and returned to standard bedding conditions. Pups were weighed and their body weight was recorded on PND 11, 28, 60, and 110 to assess development.

**Behavior:** Behavioral testing was conducted during the light phase. Mice were moved to the testing room and habituated for at least 30 minutes before experimentation. Behavioral apparatuses were cleaned with 70% ethanol between subjects. When semiautomatic video analysis was used, results were checked manually and found to be accurate. For the light-dark box (LDB), open field (OF), and ethanol-induced locomotion tasks, behavior was video-recorded (Panasonic WV-CP304) and analyzed via the CleverSys TopScan behavior analysis system. For the elevated zero maze (EZM) and repeated EZM tasks, animal behavior was video-recorded (Basler acA1300-60gm GigE) and analyzed via Noldus EthoVision XT 17 software.

**Light-Dark Box:** The LDB apparatus consisted of a novel 44.5 x 45.5 x 39 cm polycarbonate box (Stoelting Co., Wood Dale, IL, USA) evenly divided into open (light; ~260 lux) and walled/covered (dark) zones, with a small opening between. Animals were placed in the center of the open zone and allowed to explore freely for 10 minutes. Videos were analyzed to determine the number of entries into the dark zone and time spent on each side of the box.

**Elevated Zero Maze:** The EZM apparatus consisted of a ring-shaped polycarbonate runway (50 cm inner diameter, 5 cm lane width, 50 cm off the ground) with equal amounts of space devoted to open and walled quadrants (15 cm wall height; Stoelting Co.). In the basic EZM test, mice were placed in the center of the open zone (~110 lux) and allowed to explore freely for 10 minutes. In the repeated EZM task, mice were habituated to 3x saline injections (10 mL/kg, i.p.) one week before testing. The following week, mice received an injection of either saline (10 mL/kg, i.p.) or alcohol (1.2 g/kg, 10 mL/kg, i.p.; order counterbalanced) and then were immediately placed in the center of the open zone and allowed to explore freely for 10 minutes. A subsequent test was conducted one week later with the opposite injection. Factors such as time of day and lighting were kept as consistent as possible between the first and second EZM tests. Videos were analyzed to determine the time spent in the open zone. Six mice (1 control, 5 LBN) were excluded from analysis due to off target injections or the mouse falling off the maze.

**Open Field:** The OF apparatus consisted of a novel 44.5 x 45.5 x 39 cm polycarbonate box (Stoelting Co.). Mice were placed in the center of the box and allowed to explore freely for 10 minutes. Videos were analyzed to determine the time spent in the inner vs outer 50%.

**Ethanol-Induced Locomotion:** The ethanol-induced locomotion test took place in the OF apparatuses. Mice completed this task ~7 days following completion of the repeated EZM test. Thus, mice were habituated to 4x saline injections and 1x ethanol injections. On the first day of testing, mice were placed in the center of the box and allowed to explore for 60 minutes. Following this, mice received either saline (10 mL/kg, i.p.) or ethanol (1.2 g/kg, 10 mL/kg, i.p.; counterbalanced) injections and were returned to the box for 60 minutes. The following day, the protocol was repeated with the opposite injection. Videos were analyzed to determine the distance traveled (m) per 5-minute time bin and time spent in the inner vs outer 50%.

**IntelliCage Social-Operant Drinking:** Five Iso Pads (15.24 x 25.4 cm; Braintree Scientific Co., ISO 6105, Braintree, MA, USA), two Nestlest squares (Ancare, NES3600), and four Mouse Houses (Tecniplast) were provided in each cage as bedding and enrichment. Before the start of testing (~PND 50), all mice were briefly anesthetized with isoflurane (Baxter, 10019-360-40, Deerfield, IL, USA) and subcutaneously implanted with a RFID transponder (1.25 x 7 mm; TSE) in the dorsocervical region for unique identification. Cohorts one, two, and three completed an operant alcohol-drinking task, while cohorts four and five conducted an operant water-drinking task.

*Habituation.* All sipper bottles contained water. Mice had free access to bottles.

*Water Training.* All sipper bottles contained water. Mice were trained to nosepoke on a fixed ratio (FR)-1 schedule to gain access to water bottles. Once an individual mouse reached a 60% or higher success rate, they moved to an FR-3 schedule for the remainder of the program duration. All mice achieved the FR-3 schedule within three days.

*Intermittent Alcohol.* (Cohorts one, two, and three only.) The sipper bottles in two of the four operant corners (8 sipper bottles) were replaced with 20% alcohol (95% ethanol [190 proof,

Deacon Labs] in tap water, v/v). Mice had continuous access to water on an FR-3 schedule and intermittent access to alcohol, for 24-hour periods (12:30 PM to 12:30 PM) every other day. Mice performed this phase of the task for either 16 (cohorts two and three) or 24 (cohort one) consecutive days.

*Operant Water Drinking.* (Cohorts four and five only.) All sipper bottles contained water. Mice had continuous access to water on an FR-3 schedule. Mice performed this phase of the task for 14 days. An issue with the program resulted in accidental water deprivation among cohort four mice. Due to the confounding effects of stress, data from cohort four has been excluded from all drinking analysis.

*Alcohol Adulteration.* (Cohorts one, two, and three only.) After completing Intermittent Alcohol (cohort one) or halfway between two 16-day periods of Intermittent Alcohol (cohorts two and three), the sipper bottles containing 20% alcohol were replaced with 0.5 mM, and subsequently, 1 mM quinine adulterated 20% alcohol to assess punishment-insensitive drinking. Access to sipper bottles continued on an FR-3 operant schedule. This program was run for four days total, with one day of access to 20% alcohol adulterated with 0.5 mM of quinine, one day of access to 20% alcohol adulterated with 1 mM of quinine, and two days of access to sipper bottles containing water only.

*Water Adulteration.* (Cohorts four and five only.) After completing Operant Water Drinking, the sipper bottles in two of the four operant corners were replaced with water adulterated with 0.5 mM, and subsequently, 1 mM quinine to assess baseline sensitivity to quinine. Access to sipper bottles continued on an FR-3 operant schedule. This program was run for four days total, with one day of access to unadulterated water in between the two days of 0.5 mM and 1 mM quinine adulterated water.

*Open Bar.* (Cohorts one, two, and three only.) The sipper bottles in two of the four operant corners contained unadulterated 20% alcohol. Mice had free access (FR-0) to sipper bottles (i.e., the doors in all corners were open). This phase of the experiment was run for only one day.

*Data analysis.* Data was extracted using the TSE analyzer software, exported into R, and analyzed using publicly available packages from the tidyverse. For analysis of percent reduction from baseline drinking following quinine adulteration, mice with a baseline value lower than 10 were excluded (n = 6).

**Quantitative Polymerase Chain Reaction:** Mice were deeply anesthetized with isoflurane and rapidly decapitated. Brains were extracted, chilled in 1x PBS on ice for 60 seconds, and sliced using a 1mm coronal brain matrix. DMS and NAc samples were taken using tissue punches (Miltex® Biopsy Punch with Plunger, Ted Pella; DMS: 1.5 mm, 15110-15; NAc: 1 mm, 15110-10, Redding, CA, USA). Brain punches were placed in RNAlater, homogenized, and total RNA was purified using RNeasy Plus Mini kit (QIAGEN, 74192, Redwood City, CA, USA). cDNA was synthesized using iScript Reverse Transcription Supermix (BioRad, 1708841, Hercules, CA, USA). *Actb* (Mm01205647), Drd*1* (Mm02620146_s1), and *Drd2* (Mm00438541_m1) TaqMan Gene Expression Assays (Applied Biosystems, Waltham, MA, USA) were used to determine relative mRNA expression. Samples were run in duplicate and in parallel with negative controls using the QuantStudio 3 Real-Time PCR System (Thermofisher Scientific, Waltham, MA). Plates were designed to balance the distribution of sex and rearing conditions of the samples. The cycling conditions were: initial holds at 50°C (2 min) and 95°C (10 min), 40 cycles of 95°C (15 s) and 60°C (1 min). Expression of *Drd1* and *Drd2* was calculated relative to region-specific group averages of values among alcohol-naïve control mice via the ΔΔCt method, with *Actb* as the internal control gene. Relative expression values greater than 6 (n = 2) were excluded from the analysis.

**Autoradiography:** Mice were deeply anesthetized with isoflurane and rapidly decapitated. Brains were extracted and flash-frozen in 2-methylbutane before sectioning (20 μm) on a Cryostat (CryoStar NX-50, Epredia, Kalamazoo, MI, USA). Sections were thaw-mounted on to either positively- or non-positively-charged slides, and the latter were subsequently baked at 60°C for 10 min to ensure tissue adhesion. Testing confirmed that baking the sections onto non-positively-charged slides did not affect radioligand binding. Slides were pre-incubated for 10 min at room temperature in washing buffer (50 mM Tris-HCl, pH 7.4 with 120 mM NaCl, 1 mM MgCl2, 5 mM KCl, 2 mM CaCl2) and then incubated for 60 min in the same buffer conditions containing either [$^3$H]raclopride (4 nM, 81.8 Ci/mmol, Revvity) or [$^3$H]SCH-23390 (2.5 nM, 83.9 Ci/mmol, Revvity) with Ketanserin Tartrate (40 nM, Tocris, Bristol, UK) to block off-target serotonergic binding by SCH-23390. Non-specific binding was determined by incubating a subset of slides in the same conditions in the presence of either butaclamol (10 µM, Tocris) or SCH-23390 (10 µM, Tocris). Slides were then washed 2x 15 seconds in ice-cold Tris-HCl buffer (50mM) and dipped in ice-cold distilled water to remove salts. The slides were apposed to a BAS-TR2025 Storage Phosphor Screen (Fujifilm, Kawasaki, JP) along with a Carbon-14 Standards slide (American Radiolabeled Chemicals Inc., St Louis, MO, USA) inside a Hypercassette (Amersham Biosciences, Amersham, UK) for 12 days and then imaged using a Typhoon biomolecular imager (Cytiva, Marlborough, MA, USA). The digitized images were calibrated using the Carbon-14 Standards slide and radioactivity quantified using Multigauge software (GE Healthcare, Chicago, IL, USA). Regions of interest (ROI; 6 to 10 ROI per slice) were manually drawn using neuroanatomical landmarks. Specific binding was calculated by subtracting non-specific binding (nCi/g) from each ROI, and percent binding was determined by normalizing these values to the mean specific binding in the corresponding striatal subregions of alcohol-naïve, control mice.

**Fast-Scan Cyclic Voltammetry:** Mice were deeply anesthetized with isoflurane and rapidly decapitated. Brains were extracted, mounted on a vibratome (VT-1200S, Leica Microsystems, Wetzlar, GR), and sliced coronally in a warm oxygenated cutting solution (32°C), containing the following (in mM): 90 sucrose, 80 NaCl, 24 NaHCO3, 1.25 NaH2PO4, 10 glucose, 3.5 KCl, 0.5 CaCl2, 4.5 MgCl2, and 3 kynurenic acid. Slices were incubated for 20 min at 32°C in artificial CSF (ASCF) containing the following (in mM): 124 NaCl, 1 NaH2PO4, 2.5 KCl, 1.3 MgCl2, 2.5 CaCl2, 20 glucose, 26.2 NaHCO3, and 0.4 ascorbic acid, and then maintained at room temperature until recordings. Fast-scan cyclic voltammetry (FSCV) recordings were conducted in the DMS. Slices were submerged in a chamber with continuous perfusion at 2 ml/min with ACSF heated to 32°C using an inline heater (Harvard Apparatus, Holliston, MA, USA). Cylindrical carbon-fiber electrodes were prepared with T650 fibers (7 μm diameter, ~150 μm of exposed fiber) inserted into a glass pipette. The carbon-fiber electrode was held at −0.4 V versus Ag/AgCl and a triangular voltage ramp (−0.4 to +1.2 and back to −0.4 V at 0.4 V/ms) was delivered every 100 ms. Dopamine transients were electrically evoked, a glass pipette filled with ACSF was placed near the tip of the carbon fiber (~100–200 μm), and a rectangular pulse (0.2 ms) was applied every 2 min. Data were collected with a retrofit head stage (CB-7B/EC with 5 MΩ resistor) using a Multiclamp 700B (Molecular Devices, San Jose, CA, USA) amplifier after low-pass filtering at 10 kHz and digitized at 100 kHz using a NI USB-6229 (National Instruments, San Jose, CA) I/O board. The output was recorded using Igor Pro (Wavemetrics, Portland, OR, USA) using custom acquisition software utilizing parts of mafPC (courtesy of M. A. Xu-Friedman) and analyzed by custom written software in Igor Pro. Baseline voltammograms before stimulation were averaged and subtracted from the voltammograms from the selected time window, and transients were calculated from the oxidation peak region. The current peak amplitude of the evoked dopamine transients was converted to dopamine concentration according to the post-experimental calibration of the carbon-fiber electrodes with dopamine (1–3 μM) applied locally through a glass pipette in the recording chamber. Transients were compared before and after bath application of 1 μM DHßE.

Figure S1 Anderson et al

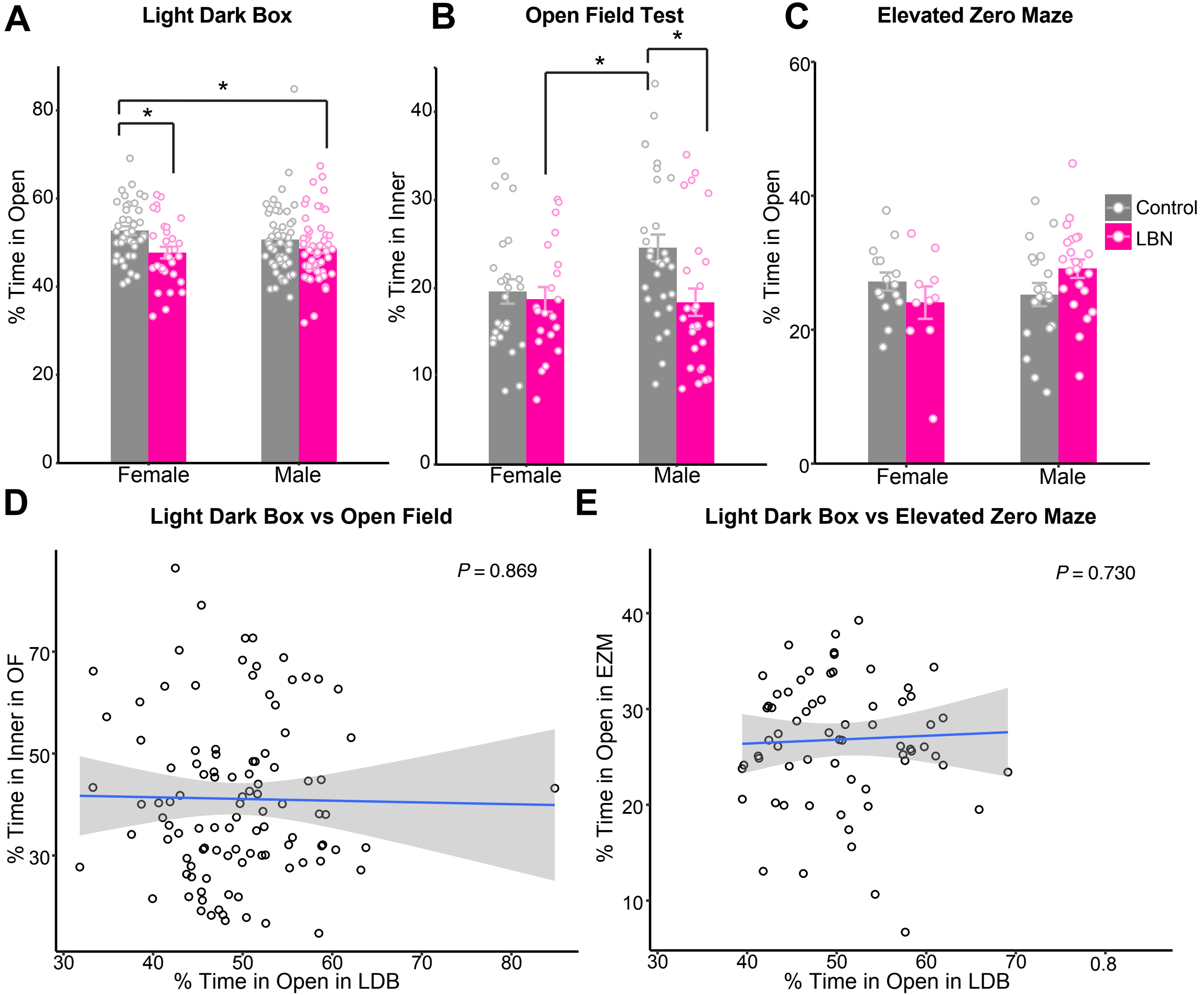

### SUPPLEMENTARY FIGURES

**Supplemental Figure 1. (A, B, C)** Bars showing mean (and SEM) percent time in open/inner zones over 10 minutes among control and LBN-reared mice in **(A)** light dark box (ANOVA, condition: $F(1) = 8.79$, $p = 0.0035$; Tukey's, F LBN vs F control: $p = 0.028$, M LBN vs F control: $p = 0.042$; F: n = 41 control, 31 LBN; M: 49 control, 54 LBN), **(B)** open field (ANOVA, $F(1) = 6.24$, $p = 0.014$; Tukey's, LBN F vs control M: $p = 0.037$, LBN M vs control M: $p = 0.015$; F: n = 26 control, 21 LBN; M: 30 control, 27 LBN), and **(C)** elevated zero maze tasks (ANOVA, condition x sex: $F(1) = 4.08$, $p = 0.048$, F: 15 control, 10 LBN; M: 19 control, 23 LBN) separated by sex. **(D)** Correlation between percent time in open in the light dark box with percent time in the inner of the open field apparatus (linear regression, $y = 42.8 - 3.41x$, $R^2_{adj} < 0.01$, $F(1, 102) = 0.027$, $p = 0.87$; n = 104). **(E)** Correlation between percent time in open in the light dark box with that in the elevated zero maze task (linear regression, $y = 24.8 + 4.07x$, $R^2_{adj} < 0.01$, $F(1, 65) = 0.12$, $p = 0.73$; n = 67). For panels **(D-E)**, points represent individual data points and shading shows 95% confidence interval. For all panels, *$p < 0.05$. Mice are color-coded by rearing condition (control: gray, LBN: pink).

Figure S2 Anderson et al

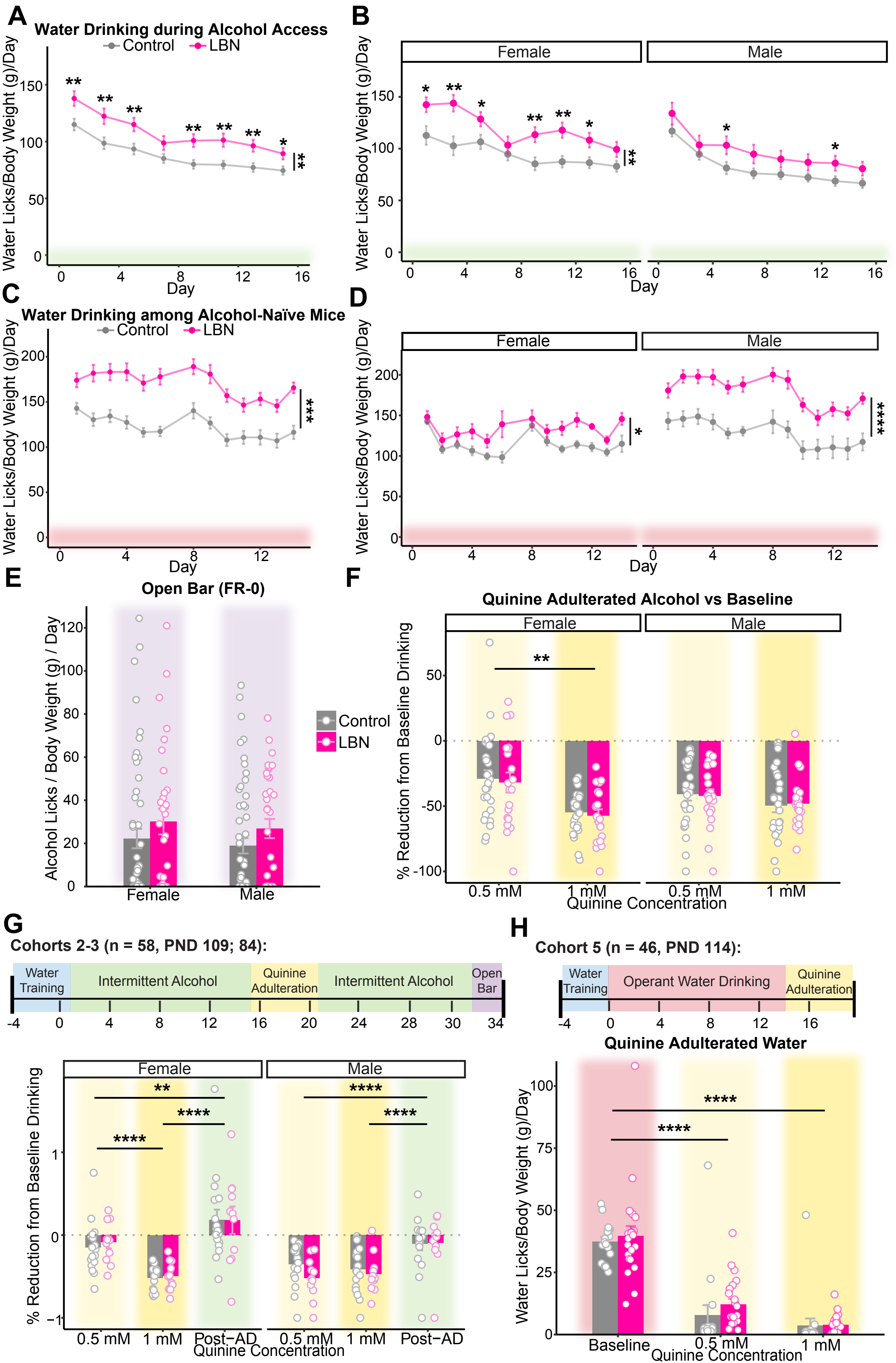

**Supplemental Figure 2. (A, B)** Water licks per day normalized by animal body weight (g; measured weekly) for 8 days of intermittent alcohol access (same period as Figure 2B-C) among alcohol-exposed (cohort 1-3) control and LBN-reared mice **(A)** collapsed by sex (Mixed ANOVA, condition: F(1, 93) = 11.71, p = 0.00093; sex: F(1, 93) = 9.01, p = 0.0030; day: F(4.26, 395.92) = 55.371, p = 8.51e-39; sex x day: F(4.26, 395.92) = 3.58, p = 0.0060; group x sex x day: F(4.26, 395.92) = 2.55, p = 0.036; t-test with Bonferroni adjustment, day 1: p = 0.0062; 3: p = 0.0061; 5: p = 0.0056; 9: p = 0.0027; 11: p = 0.0015; 13: p = 0.0035; 15: p = 0.018; n = 54 control, 43 LBN) and **(B)** separated by sex (F: left; Mixed ANOVA, condition: F(1, 44) = 7.86, p = 0.0070; day: F(4.2, 184.93) = 22.81, p = 7.54e-16; condition x day: F(4.2, 184.93) = 3.03, p = 0.017; t-test with Bonferroni adjustment, day 1: p = 0.019; 3: p = 0.0023; 5: p = 0.034; 9: p = 0.0057; 11: p = 0.0018; 13: p = 0.017; n = 26 control, 20 LBN; male: right; Mixed ANOVA, day: F(3.82, 187.09) = 36.70, p = 5.92e-22; t-test with Bonferroni adjustment, day 5: p = 0.039; 13: p = 0.046; n = 28 control, 23 LBN). **(C, D)** Water licks per day normalized by animal body weight (g; measured weekly) for days 1-14 of operant water drinking (except day 7 due to a cage change) among alcohol-naïve (cohort 5) control and LBN-reared mice **(C)** collapsed by sex (Mixed ANOVA, condition: F(1, 42) = 15.91, p = 0.00026; sex: F(1, 42) = 11.24, p = 0.0020; day: F(3.81, 160.05) = 9.09, p = 1.89e-06; sex x day: F(3.81, 160.05) = 7.71, p = 1.51e-05; t-test with Bonferroni adjustment, day 1: p = 0.0034; 2: p = 8.81E-05; 3: p = 0.00013; 4: p = 2.76E-05; 5: p = 2.51E-06; 6: p = 9.65E-07; 8: p = 2.06E-04; 9: p = 7.89E-05; 10: p = 8.89E-06; 11: p = 0.0014; 12: p = 0.00025; 13: p = 0.00057; 14: p = 4.78E-06; n = 22 control, 24 LBN) and **(D)** separated by sex (F: left; Mixed ANOVA, condition: F(1, 12) = 16.59, p = 0.0020; day: F(3.69, 44.33) = 5.55, p = 0.0010; t-test with Bonferroni adjustment, day 4: p = 0.035; 5: p = 0.040; 6: p = 0.019; 10: p = 0.0099; 11: p = 0.0047; 12: p = 0.0058; n = 9 control, 5 LBN; M: right; Mixed ANOVA, condition: F(1, 30) = 20.61, p = 8.52e-05; day: F(3.13, 93.95) = 17.70, p = 2.02e-09; t-test with Bonferroni adjustment, day 1: p = 0.011; 2: p = 0.00024; 3: p = 0.00063; 4: p = p = 0.00036; 5: p = 1.20E-05; 6: p = 4.25E-05; 8: p = 0.00075; 9: p = 0.00053; 10: p = 0.00028; 11: p = 0.016; 12: p = 0.0042; 13: p = 0.0050; 14: p

= 0.00014; n = 11 control, 19 LBN). For panels **(A-D)**, points show mean and bars show SEM. **(E)** Alcohol licks per gram of body weight (g) among alcohol-exposed (cohort 1-3) mice during non-operant (FR-0) drinking (n.s.; n = 54 control, 43 LBN). **(F)** Percent reduction in baseline drinking in response to quinine (0.5, 1 mM) adulteration ($\% \text{ Reduction from Baseline} = 100 - \frac{\text{Alcohol Licks/Body Weight (g)/Day}}{\text{Pre-adulteration Baseline}}$) among female (left) and male (right) alcohol-exposed (cohort 1-3) mice (mixed ANOVA, day: F(1.78, 162.19) = 206.102, p = 1.27e-42; sex x day: F(1.78, 162.19) = 6.01, p = 0.0040; t-test with Bonferroni adjustment, F: p = 0.0030; F: n = 26 control, 20 LBN; M: 28 control, 23 LBN). **(G)** Experimental timeline Percent reduction in baseline drinking during and after quinine adulteration (0.5 mM, 1 mM) among female (left) and male (right) alcohol-exposed (cohort 2-3) mice (mixed ANOVA, sex: F(1, 52) = 6.78, p = 0.012; day: F(2.07, 107.56) = 54.39, p = 2.18e-17; sex x day: F(2.07, 107.56) = 8.67, p = 0.00028; t-test with Bonferroni adjustment, F(0.5 vs 1 mM): p = 4.50E-05; F(0.5 mM vs post-AD): p = 0.0018; F(1 mM vs post-AD): p = 9.25E-13; M(0.5 mM vs post-AD): p = 2.91E-05; M(1 mM vs post-AD): p = 8.66E-06; F: 17 control, 11 LBN; M: 18 control, 12 LBN). **(H)** Experimental timeline. Water licks per day normalized by body weight (g) prior to and during quinine adulteration (0.5 mM, 1 mM) among alcohol-naïve (cohort 5) mice (mixed ANOVA, day: F(1.2, 42.15) = 94.56, p = 2.36e-13; t-test with Bonferroni adjustment: 0.5 mM vs baseline: p = 5.17E-17; 1 mM vs baseline: p = 2.05E-22). For panels **(E-H)**, bars showing mean and SEM are overlaid with data from individual mice. For all panels, *p <0.05, **p < 0.01, ***p<0.001, ****p < 0.0001. Mice are color-coded by rearing condition (control: gray, LBN: pink).

Figure S3 Anderson et al

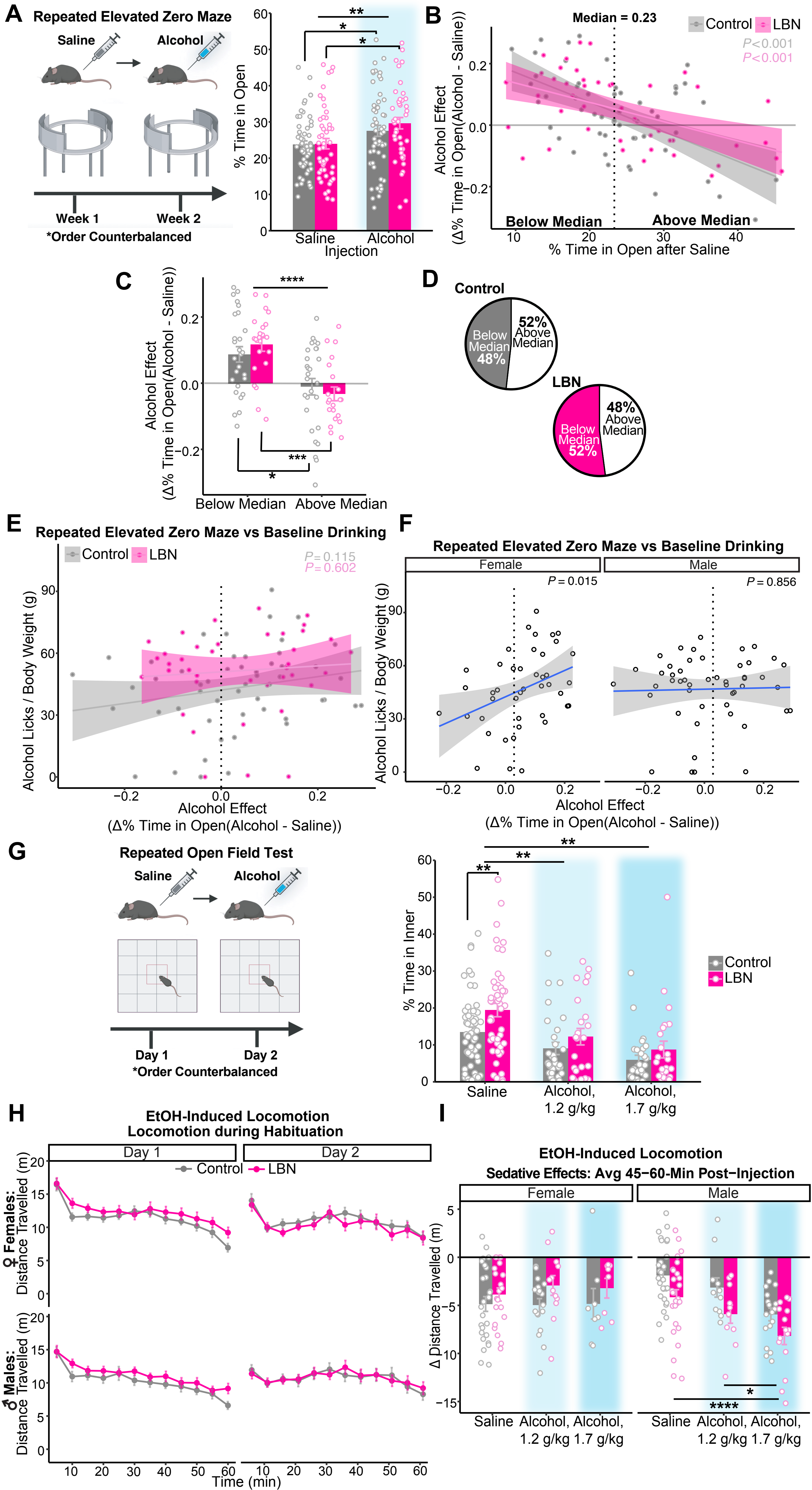

**Supplemental Figure 3. (A)** Experimental design. Percent time in open among control and LBN-reared mice after saline vs alcohol (1.2 g/kg, i.p.; order counterbalanced) injection (mixed ANOVA, injection: F(1, 96) = 11.05, p = 0.04; Tukey's, control: p = 0.041; LBN: p = 0.0091; n = 55 control, 44 LBN). Bars showing mean and SEM are overlaid with data from individual mice. **(B)** EZM performance after alcohol vs saline injection was compared for each mouse: $\text{Alcohol Effect} = \text{\% Time in Open}_{\text{Alcohol}}\text{- \% Time in Open}_{\text{Saline}}$. Correlation between percent time in open after saline injection and alcohol effect among control (linear regression, y = 26.7 - 96.4x, $R^2_{adj}$= 0.29, F(1,53) = 23.2, p = 1.26e-05, n = 55) and LBN-reared (y = 19.9 - 62.8x, $R^2_{adj}$= 0.25, F(1,42) = 15.2, p = 0.00035, n = 44) mice, with symbols representing individual data points and shading showing 95% confidence interval. Dashed line shows median value used to split population into below (% time in open < 23.2) and above median (% time in open > 23.2) subgroups. **(C)** Alcohol effect among below vs above median subgroups (ANOVA, F(1) = 27.93, p = 8.01E-07; Tukey's, control: p = 0.010; LBN: p = 0.00020; below median: n = 27 control, 25 LBN; above median: n = 29 control, 23 LBN). Bars showing mean and SEM are overlaid with data from individual mice. **(D)** Within control (top) and LBN (bottom) groups, the percentage of mice within above vs below median subgroups (control: 51.79% above, 48.21% below; n = 56; LBN: 47.92% above, 52.08% below; n = 48). **(E, F)** Correlation between alcohol effect and baseline drinking **(E)** collapsed by sex (control: linear regression, y = 42.1 + 32.5x, $R^2_{adj}$ = 0.03, F(1, 51) = 2.58, p = 0.115; n = 53; LBN: y = 51 + 14.5x, $R^2_{adj}$ < 0.01, F(1, 37) = 0.28, p = 0.60; n = 39) and **(F)** separated by sex (F: left; linear regression, y = 42.5 + 74.3x, $R^2_{adj}$ = 0.11, F(1, 41) = 6.41, p = 0.015; n = 43; M: right; y = 46.8 + 3.6x, $R^2_{adj}$ < 0.01, F(1, 47) = 0.033, p = 0.856; n = 49). Points represent individual data points and shading shows 95% confidence interval. **(G)** Experimental timeline. Percent time in inner among control and LBN-reared mice after saline vs alcohol (1.2 g/kg: n = 29 control, 24 LBN; 1.7 g/kg: n = 27 control, 24 LBN; Mixed ANOVA, Alcohol, 1.2 g/kg vs saline: F(1, 51) = 10.06, p = 0.0030; Alcohol 1.7 g/kg vs saline: F(1, 46) = 18.08, p = 0.00010; t-test w/ Bonferroni adjustment: saline: $p = 0.0090$). Bars showing mean and SEM are overlaid with data from individual mice. Cartoon is made from modified BioRender templates (license Anderson, L. (2025) https://BioRender.com/wzzh1vz). **(H)** Distance travelled (m) per 5-min time bin during habituation in the ethanol-induced locomotion task among female (left; ANOVA, n.s.; n = 26 control, 21 LBN) and male mice (right; ANOVA, n.s.; n = 30 control, 26 LBN). Points show mean and bars show SEM. **(I)** Averaging difference in distance travelled 45-60 minutes post-injection among female (left; ANOVA, n.s.; saline: n = 26 control, 21 LBN; alcohol, 1.2 g/kg: n = 18 control, 12 LBN; alcohol, 1.7 g/kg: 8 control, 8 LBN) and male mice (right; ANOVA, condition: $F(1) = 12.97$, $p = 0.00049$; injection: $F(2) = 14.61$, $p = 2.52e-06$; Tukey's, alcohol, 1.7 g/kg vs saline: $p = 1.22E-06$; alcohol, 1.2 g/kg vs alcohol, 1.7 g/kg: $p = 0.022$; n = 30 control, 26 LBN; alcohol, 1.2 g/kg: n = 11 control, 11 LBN; alcohol, 1.7 g/kg: n = 18 control, 15 LBN). Bars showing mean and SEM are overlaid with data from individual mice. For all panels, *$p <0.05$, **$p < 0.01$, ***$p<0.001$, ****$p < 0.0001$. Mice are color-coded by rearing condition (control: gray, LBN: pink).

Figure S4 Anderson et al

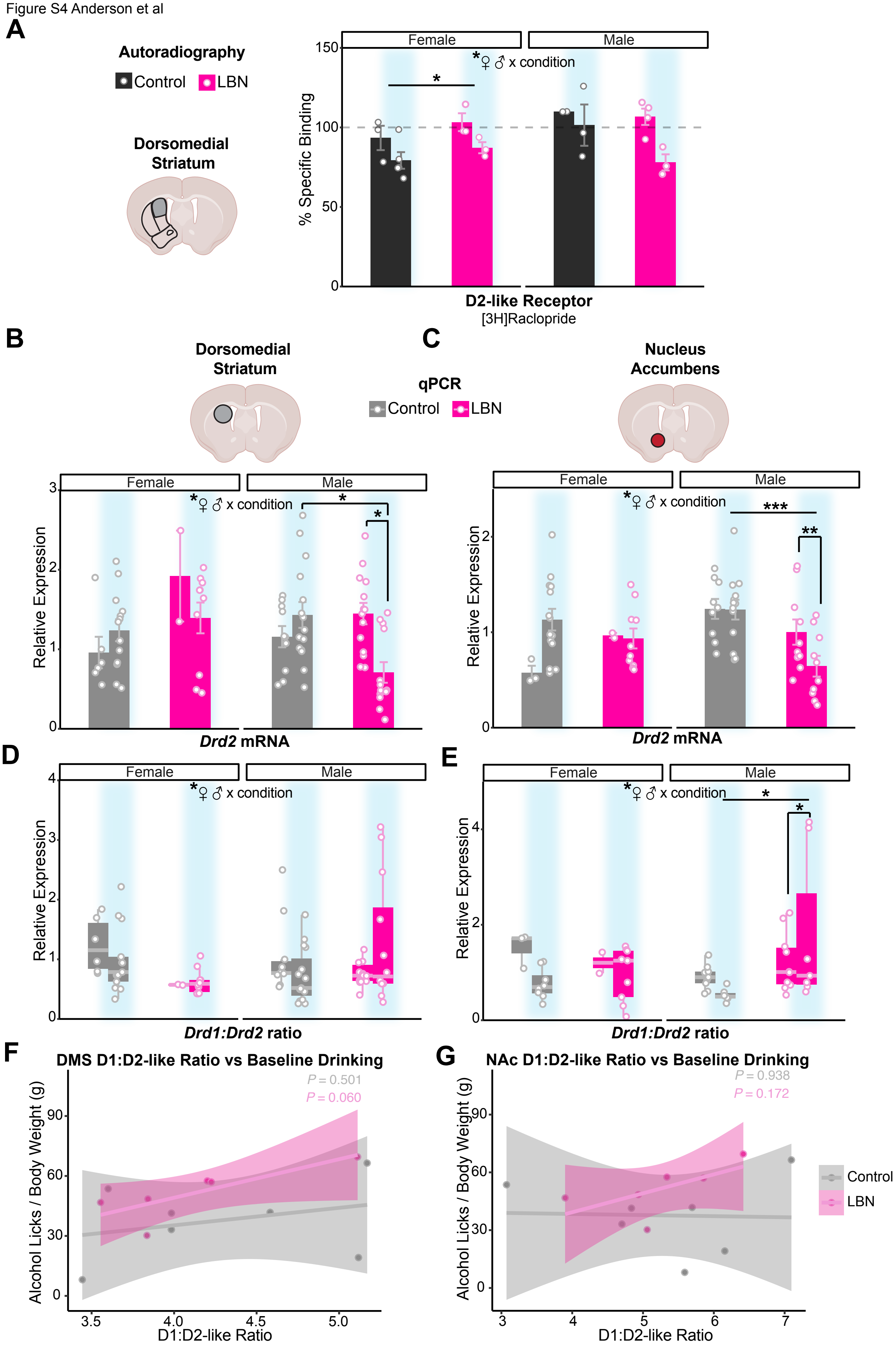

**Supplemental Figure 4. (A)** Percent specific binding of [3H]Raclopride in the DMS of alcohol-naïve and -exposed (blue highlight) control (black) and LBN-reared (pink) mice, split by sex (F: left; control: n = 3 naïve, 5 exposed; LBN: n = 3 naïve, 3 exposed; M: right; control: n = 2 naïve, 3 exposed; LBN: 4 naïve, 3 exposed; ANOVA, alcohol: F(1) = 13.39, p = 0.0018; condition x sex: F(1) = 5.42, p = 0.032; Tukey's, LBN(exposed vs naïve): p = 0.018; control (F vs M): p = 0.042). **(B, C)** Relative expression of *Drd2* among alcohol-naïve and -exposed (blue highlight) control (gray) and LBN-reared (pink) mice split by sex in the **(B)** DMS (F: left; control: n = 6 naïve, 13 exposed; LBN: n = 2 naïve, 10 exposed; M: right; control: n = 10 naïve, 15 exposed; LBN: n = 14 naïve, 13 exposed; ANOVA, condition x alcohol: F(1) = 10.71, p = 0.0016; condition x sex: F(1) = 10.68, p = 0.0016; Tukey's, exposed M(control vs LBN): p = 0.013; LBN M(exposed vs naïve): p = 0.012; LBN(F vs M): p = 0.027; LBN(exposed vs naïve): p = 0.21) and **(C)** NAc (F: left; control: n = 3 naïve, 14 exposed; LBN: n = 2 naïve, 10 exposed; M: right; control: n = 9 naïve, 13 exposed; LBN: 10 naïve, 10 exposed; ANOVA, condition: F(1) = 10.51, p = 0.0019; condition x sex: F(1) = 5.90, p = 0.018; alcohol x sex: F(1) = 4.67, p = 0.035; Tukey's, M(control vs LBN): p = 0.00094; exposed M(control vs LBN): p = 0.0065). **(D, E)** Ratio of the relative expression of *Drd1:Drd2* among alcohol-naïve and -exposed (blue highlight) control (gray) and LBN-reared (pink) mice split by sex in the **(D)** DMS (F: left; control: n = 6 naïve, 13 exposed; LBN: n = 2 naïve, 10 exposed; M: right; control: n = 10 naïve, 15 exposed; LBN: n = 14 naïve, 12 exposed; ANOVA: condition x alcohol: F(1) = 3.99, p = 0.049; condition x sex: F(1) = 7.90, p = 0.0063; Tukey's, n.s.) and **(E)** NAc (F: left; control: n = 3 naïve, 14 exposed; LBN: n = 2 naïve, 10 exposed; M: right; control: n = 9 naïve, 13 exposed; LBN: 10 naïve, 10 exposed; ANOVA, condition: F(1) = 5.02, p = 0.030; condition x sex: F(1) = 5.30, p = 0.0260; Tukey's, M(control vs LBN): p = 0.018; exposed M(control vs LBN): p = 0.045). For panels **(A-E)**, mean (bars) and SEM (error bars) are overlaid with data from individual mice. Correlation between the ratio of [3H]SCH-23390:[3H]Raclopride binding and baseline drinking in the Intellicages in the **(F)** DMS (control: linear regression, y = 0.357 + 8.74x, $R^2_{adj}$ < 0.01, F(1, 5) = 0.525, p = 0.501; n = 7; LBN: y = 28 + 19.3x, $R^2_{adj}$ = 0.54, F(1, 4) = 6.76, p

= 0.060; n = 6) and **(G)** NAc (control: linear regression, y = 40.7 - 0.57x, $R^2_{adj}$ < 0.01, F(1, 5) = 0.00674, p = 0.938; n = 7; LBN: y = -0.0154 + 9.83x, $R^2_{adj}$ = 0.26, F(1, 4) = 2.75, p = 0.172; n = 6). For panels **(F-G)**, points represent individual data points and shading shows 95% confidence interval. For all panels, *p <0.05, **p < 0.01, ***p<0.001. Cartoons in panels **(A-C)** are made from modified BioRender templates (license Anderson, L. (2025) https://BioRender.com/wzzh1vz).
